# Regenerative growth is constrained by *brain tumor* to ensure proper patterning in *Drosophila*

**DOI:** 10.1101/615948

**Authors:** Syeda Nayab Fatima Abidi, Felicity Ting-Yu Hsu, Rachel K. Smith-Bolton

**Affiliations:** Department of Cell and Developmental Biology, University of Illinois at Urbana-Champaign, Urbana, IL 61801, USA; Carle R. Woese Institute for Genomic Biology, University of Illinois at Urbana-Champaign, Urbana, IL, 61801, USA

## Abstract

Some animals respond to injury by inducing new growth to regenerate the lost structures. This regenerative growth must be carefully controlled and constrained to prevent overgrowth and to allow correct organization of the regenerating tissue. However, the factors that restrict regenerative growth have not been identified. Using a genetic ablation system in the *Drosophila* wing imaginal disc, we have identified one mechanism that constrains regenerative growth, impairment of which also leads to erroneous patterning of the final appendage. Regenerating discs with reduced levels of the RNA-regulator Brain tumor (Brat) exhibit enhanced regeneration, but produce adult wings with disrupted margins that are missing extensive tracts of sensory bristles. In these mutants, aberrantly high expression of the pro-growth factor Myc and its downstream targets likely contributes to this loss of cell-fate specification. Thus, Brat constrains the expression of pro-regeneration genes and ensures that the regenerating tissue forms the proper final structure.

**Author Summary:** While much has been published about the signals that stimulate regeneration, the mechanisms that constrain and/or terminate regeneration have not been well characterized. Thus, we do not understand what limits may exist on the rate of regenerative growth, what mechanisms constrain regeneration, and what the consequences might be of enhancing regrowth. Here, we detail our discovery and characterization of a mechanism that constrains regeneration, and the deleterious effects of reducing that constraint. In this manuscript, we describe our identification of the RNA regulator Brat as a factor that constrains regenerative growth. Without this constraint on regenerative growth, patterning mistakes occur leading to a malformed regenerated structure. We demonstrate that the patterning errors are not due to faster growth itself, but are due to the overexpression of the pro-growth transcription factor Myc.

## Introduction

Regeneration is the remarkable process by which some organisms replace tissues and organs after damage such that both morphology and function are restored. Complete regeneration requires several steps to occur correctly including wound healing, cell proliferation, and proper patterning and cell-fate specification in the newly formed tissue. The degree of regenerative capacity varies among different species, ranging from whole-body regeneration in hydra and planaria to limited tissue regeneration in mammals. Work in several model organisms has identified signaling pathways and molecular mechanisms that are important for initiating and executing regenerative growth after tissue damage, including JNK signaling (1–5), JAK/STAT signaling (6–8), EGFR signaling (9–12), Hippo signaling (13–17), Wnt signaling (18–24), and Myc (23,25). Many of these mechanisms are also important during normal development, and the process of regeneration was traditionally thought to be a redeployment of earlier developmental steps (3,9,25–29). However, recent evidence suggests that regeneration is not a simple reiteration of development but can employ regeneration-specific regulatory mechanisms (3,25,30–34). Indeed, faithful regeneration likely requires additional mechanisms, since regrowth happens in the presence of wound-response signaling and in a developed juvenile or adult organism. Additionally, pro-growth pathways that are used during normal development are often activated in new ways and at higher strengths in the regenerating tissue (2,7,15,23). These augmented pro-growth signals must be constrained as regeneration progresses to prevent overgrowth and to enable re-establishment of pattern and cell-fate specification. Thus, growth suppressors and additional patterning factors are likely used to terminate regeneration and allow differentiation (35). However, despite our understanding of the pro-growth signals needed for regeneration, we do not yet know what factors exist in different model organisms to restrain growth and promote re-patterning of regenerating tissue.

*Drosophila melanogaster* imaginal discs, precursors of adult fly appendages, are simple columnar epithelia that have well-characterized, complex expression of patterning genes that determine cell-fate specification. Imaginal discs undergo regeneration after damage (36), and we have previously used a genetic ablation system to study patterning in the regenerating tissue (23,32). Here we identify the RNA-regulator Brain tumor (Brat) as a key constraint on regenerative growth that ensures proper formation of the regenerated structure. Brat is a member of the TRIM- (tripartite motif containing)- NHL (NCL-1, HT2A, and LIN-41) family of proteins and functions as a translational repressor by binding to its target RNAs either independently or in a complex with Pumilio and Nanos (37–39). It acts as a potent differentiation factor and tumor suppressor in neural and ovarian germline stem cell lineages (40–43). Human and mouse orthologs of Brat, TRIM3 and TRIM32 respectively, also possess tumor-suppressor activity in glioblastomas and are required for neuronal and muscle differentiation (44–47).

We show that regenerating wing imaginal discs with reduced levels of Brat regenerate better than controls, but the resulting adult wings have a disrupted margin. The margin loses some of the characteristic sensory bristles and veins, demonstrating an error in cell-fate specification. Importantly, these phenotypes are regeneration-specific, as they are not observed in the mutant animals after normal development. The enhanced regeneration is due to increased expression of the growth regulators Myc and Wingless as well as upregulation of *ilp8,* which delays metamorphosis and allows the damaged tissue more time to regenerate. Intriguingly, this seemingly beneficial elevated Myc expression can also cause aberrant cell-fate specification at the wing margin. This disruption of patterning does not occur through general enhanced proliferation, but may be due to misregulation of Myc target genes, including Chronologically inappropriate morphogenesis (Chinmo). Hence, Brat acts as an important growth regulator and protective factor during regeneration, by constraining levels of Myc, Chinmo, and possibly other Myc targets to prevent errors in patterning, cell-fate specification, and differentiation in the regenerating tissue.

## Results

### Brat suppresses regenerative growth and is required for wing margin cell-fate specification during regeneration

To identify genes important for regenerative growth and re-patterning, we performed a candidate screen, using our wing imaginal disc ablation system (23). The primordial wing was targeted for ablation at the early third-instar larval stage by using *rotund-GAL4* to drive the expression of the proapoptotic gene *reaper* for 24 hours (Figure 1A). Our ability to restrict damage to 24 hours was provided by *tubulin-GAL80^ts^*, which can inhibit GAL4 activity at 18°C, but allows GAL4-driven cell death at 30°C in the 24-hour window. The extent of wing imaginal disc regeneration in the larvae was reflected in the adult wing size. Hence, the resulting adult wings were scored based on size and patterning features to identify mutants that affect genes that are involved in regulating regenerative growth and establishment of cell fates. There is inherent variability in this system because of its sensitivity to environmental conditions such as temperature, humidity, and food quality, causing the results of different experiments to vary slightly (14,48–51). Animals with the same genotype within an experiment also showed some variation, due to stochastic differences in the time each animal takes to eclose, with animals that take longer to eclose having larger wings (23,50). However, differences between control and mutant animals using this system are reproducible, consistent, and have identified key regeneration genes (32,49–51).

**Figure 1.**
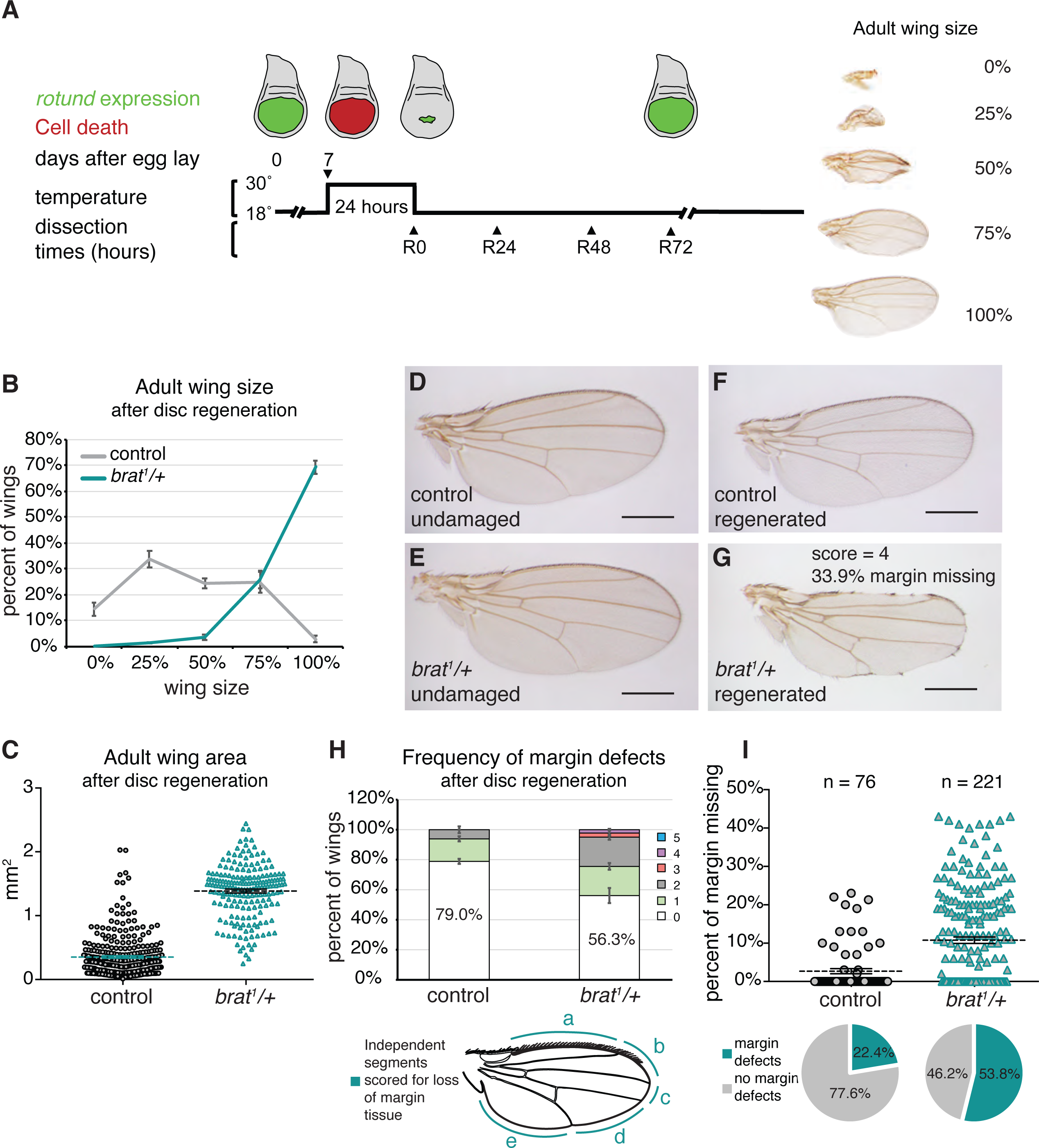
Enhanced regenerative growth and wing margin cell-fate specification defects in *brat^1^/+* during regeneration. (A) The protocol used to study regeneration. Animals were raised at 18°C and shifted to 30°C for 24 hours during early third-instar larval development on day 7 after egg lay (AEL). Larvae were returned to 18°C and were dissected at the time points noted during recovery (R) or allowed to pupariate and eclose. Representative wings depicting the range of adult wing sizes observed after regeneration compared to the size of a normal wing are shown. (B) Adult wing sizes observed after disc regeneration for control (*w^1118^*) (n = 317) and *brat^1^/+* (n = 208) wings, from three independent experiments. (C) Adult wing area after disc regeneration, measured using ImageJ after mounting and imaging wings, for control (*w^1118^*) (n = 309) and *brat^1^/+* (n = 195) wings. p = 2.5158E-119. Wings in (C) are from the same experiments as (B). Note that number of wings in (C) is less for both control and *brat^1^/+* due to some wings being damaged during the mounting process. (D) Undamaged control (*w^1118^*) wing. (E) Undamaged *brat^1^/+* wing. (F) Adult control (*w^1118^*) wing after disc regeneration. (G) Adult *brat^1^/+* wing after disc regeneration. (H) Frequency of margin defects seen in adult wings after disc regeneration for control (*w^1118^*) (n = 93) and *brat^1^/+* (n = 218) wings, from three independent experiments. The wing margin was divided into five segments based on where the veins intersect the margin as shown in the diagram. Each wing was scored for the number of segments that had some margin tissue missing, with wings with a perfectly intact margin scoring at zero. Wing shown in (G) had tissue missing in four segments. (I) Margin tissue lost as a percentage of total wing perimeter for control (*w^1118^*) (n = 76) and *brat^1^/+* (n = 221) wings. p = 9.947E-08. The margin perimeter and the length of margin tissue lost were measured using ImageJ after mounting and imaging wings. Wings in (I) are from the same experiments as (H). Note that number of wings in the two quantifications is different because we did not quantify wings with length <1.1 mm for males and <1.7 mm for females, to ensure analysis was being carried out on nearly fully regenerated wings. (I). Percentage of wings with no defects fell from 79.0% to 77.6% for control and from 56.3% to 53.8% for *brat^1^/+* wings due to the increased ability to detect lost margin tissue at the higher magnification and resolution achieved by imaging the wings. Wing shown in (G) had 33.9% of margin tissue missing. Error bars mark standard error of the mean (SEM). Student’s T-test used for statistical analyses. Scale bars are 0.5 mm.

Using this genetic ablation system, we identified the gene *brain tumor (brat)* as an important regulator of regenerative growth. *brat^1^/+* mutants that did not experience damage during development had adult wings that were not significantly different in size from controls (Figure S1A). However, after ablation and regeneration were induced, *brat^1^/+* mutants showed enhanced regeneration and had adult wings that were, on average, much larger than controls that had also undergone regeneration (Figure 1B and 1C). We confirmed this enhanced regeneration phenotype in heterozygotes for three other *brat* mutant alleles: *brat^192^*, *brat^150^* (52) and *brat^11^* (53)(Figure S1B).

Interestingly, we also discovered a role for *brat* in cell-fate specification during regeneration. After normal development, *brat^1^/+* mutants had adult wings that were patterned normally (Figure 1D-E and Figure S1C). To confirm that reduction in *brat* expression does not cause patterning errors during normal development, we knocked down Brat levels in the entire wing pouch using *brat* RNAi, which resulted in adult wings that were patterned normally (Figure S1D and S1E). A previous study in which Brat levels were reduced in the anterior and posterior compartments of the wing also did not report any patterning defects (54). However, when discs were ablated and allowed to regenerate, *brat* heterozygous mutant wings showed aberrant patterning such that the wing margin lost sensory bristles and vein material (Figure 1F and 1G). By contrast, control regenerated wings lost margin tissue at a lower frequency (Figure 1H and 1I). Furthermore, the extent of margin tissue lost was not as severe in control regenerated wings as compared to *brat^1^/+* regenerated wings (Figure 1H and 1I). Similar to the enhanced regeneration seen in *brat* mutants, we confirmed the loss-of-margin defect in heterozygotes for the additional three mutant alleles (Figure S1F).

### *brat* regulates entry into metamorphosis

Tissue damage in imaginal discs can induce a systemic response in the larvae, which extends the larval phase of development and delays pupariation (23,55). This delay in pupariation is due to expression of the relaxin-like peptide *ilp8* in damaged discs (56,57). To determine whether *brat* mutants regenerated better due to an enhanced delay in pupariation, we measured rates of pupariation in control and mutant animals. We found that during normal development, control and *brat^1^/+* animals pupariated at the same time, indicating that the two genotypes develop at similar rates (Figure S2A). After disc damage, *brat* mutants delayed pupariation an additional day compared to controls in which discs were also damaged (Figure 2A and Figure S2B). Note that direct comparisons cannot be made between regenerating larvae that spent 24 hours at 30°C (Figure 2A, Figure S2B) and normally developing larvae that remain at 18°C (Figure S2A), due to the effects of temperature on development. Our data show that *brat/+* mutants are able to stay in the larval stage even longer than controls, giving them more time to regenerate.

To determine why discs with reduced Brat had an increased delay in pupariation, we measured *ilp8* transcript levels through qPCR. Undamaged control animals express very low *ilp8* levels. However, after regeneration was induced, we saw an 80-fold increase in *ilp8* levels in controls, while the *brat^1^/+* animals showed a 140-fold increase (Figure 2B). Thus, *brat* suppresses *ilp8* during regeneration, regulating the timing of pupariation.

### *brat* restricts growth and proliferation during regeneration

Regenerative growth occurs through localized cell proliferation at the wound site (23,58).The proliferating cells, known as the blastema, give rise to the regenerated tissue. The blastema and the subsequent regenerated wing pouch can be labeled with the wing primordium marker Nubbin (Nub) (59). To determine whether *brat^1^/+* discs regenerated better due to increased growth rates in the wing pouch, we measured the area of the Nub-expressing cells in control and *brat^1^/+* regenerating discs. In the initial stages of regeneration, the control and mutant had similar Nub-expressing areas, indicating equal ablation and equal early regrowth. However, by 48 hours after tissue damage (recovery time 48, or R48), *brat^1^/+* wing discs had a significantly bigger Nub-expressing pouch than the control (Figure 2D-F), indicating that *brat/+* mutants were regenerating faster than controls. To assess whether this difference in growth rates was due to differences in proliferation, we counted cells going through mitosis by quantifying Phospho-histone H3 (PH3)-positive nuclei in the regenerating blastema. Reduction of *brat* resulted in a significantly higher number of PH3-positive nuclei per area at R0, but this increased proliferation had subsided to normal levels by R24 (Figure 2G-I). To confirm that the increased number of PH3-positive nuclei were not simply due to an increased number of total cells in the blastema, we also counted total Nubbin-positive nuclei in the regenerating blastema to calculate a ratio of proliferating cells to blastema cells (Figure S2C). These data confirm increased proliferation in *brat^1^/+* regenerating discs at R0. Differences in proliferation early in regeneration often become evident later when measuring wing pouch area (51). Therefore, reduction of *brat* gives the regenerating tissue a growth advantage early in regeneration, resulting in a measurable difference in tissue area by R48.

**Figure 2.**
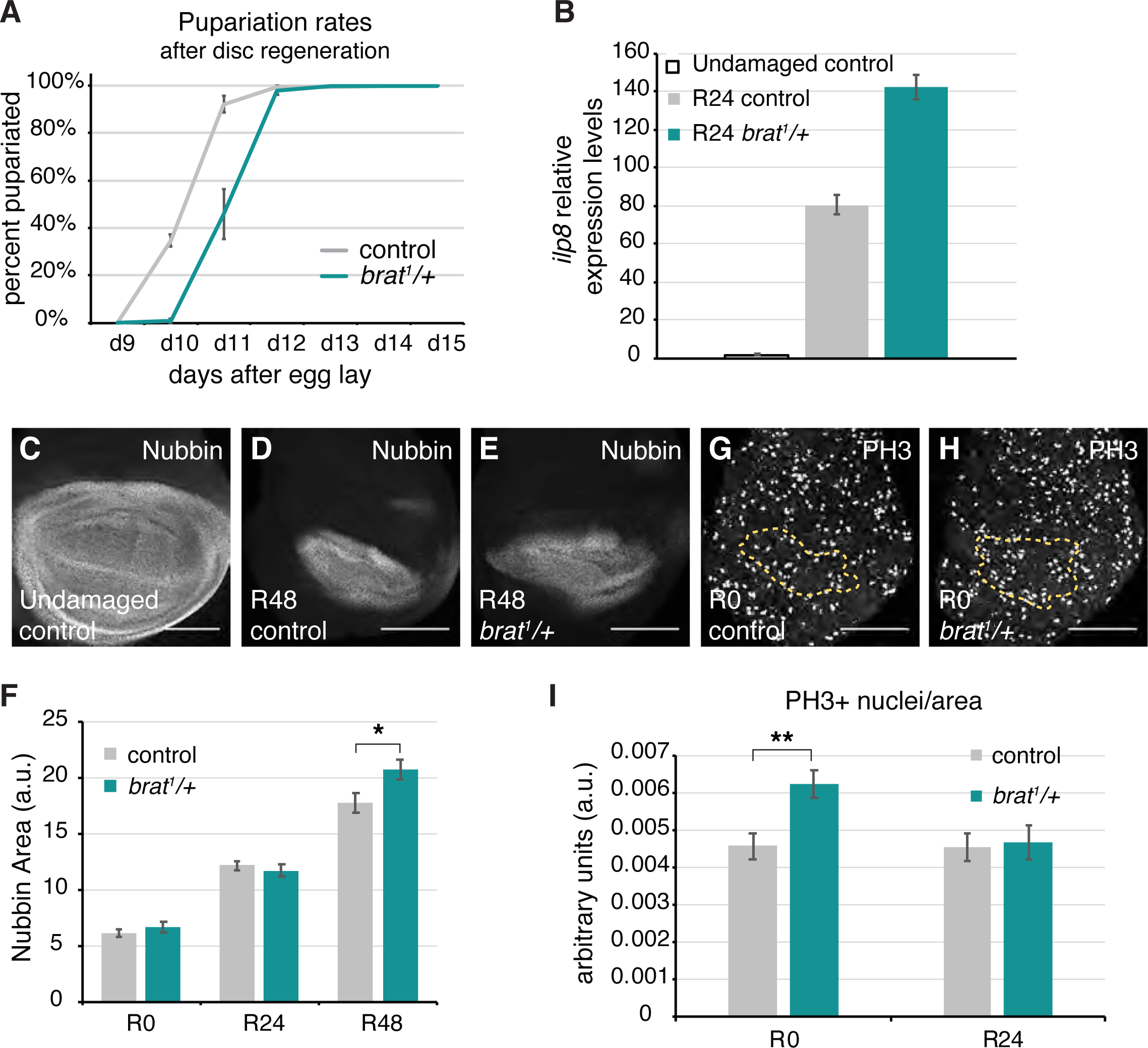
*brat^1^/+* animals have a regenerative growth advantage. (A) Pupariation rates after disc regeneration for control (*w^1118^*) (n = 384) and *brat^1^/+* (n = 107) animals, from three independent experiments. (B) Relative expression levels of *ilp8* for undamaged control, R24 control (*w^1118^*) and R24 *brat^1^/+* discs. (C) Anti-Nubbin immunostaining in an undamaged control disc. (D-E) Anti-Nubbin immunostaining in an R48 control (*w^1118^*) disc (D), and an R48 *brat^1^/+* disc (E). (F) Quantification of area of Nubbin-expressing cells for control (*w^1118^*) and *brat^1^/+* discs at R0 (n = 10 and 10), R24 (n = 12 and 12) and R48 (n = 10 and 10). * p < 0.03. (G-H) Anti-PH3 immunostaining in an R0 control (*w^1118^*) disc (G), and an R0 *brat^1^/+* disc (H). The yellow dashed lines outline the Nubbin-expressing wing pouch. (I) PH3-positive nuclei were counted within the regenerating tissue as marked by Anti-Nubbin co-immunostaining. Quantification of PH3-positive nuclei in Nubbin area for control (*w^1118^*) and *brat^1^/+* discs at R0 (n = 16 and 18) and R24 (n = 15 and 16). ** p < 0.002. Error bars represent SEM. Student’s T-test used for statistical analyses. Scale bars are 100 μm.

Wingless (Wg) and Myc are regulators of regenerative growth and are upregulated at the wound site after damage (20,21,23). Interestingly, Brat regulates stem cell differentiation in the brain by suppressing self-renewal signaling pathways such as Wnt signaling, and acting as a post-transcriptional inhibitor of Myc, to enable specification of progenitor cell fate (42,60). Additionally, Brat overexpression can suppress Myc at the post-transcriptional level in wing disc epithelial cells, although loss of *brat* does not lead to elevated Myc protein in wing discs during normal development (54). To determine whether these regulators of regenerative growth are upregulated in *brat^1^/+* regenerating discs, we examined the expression of Wg and Myc. Wg is normally expressed along the Dorso-ventral (DV) boundary and in two concentric circles at the inner and outer edge of the wing pouch (61) (Figure 3A), and Myc is expressed in the wing pouch, but is repressed in the cells at the DV boundary as they undergo cell cycle and growth arrest (62) (Figure 3B). Both Wg and Myc expression were comparable to controls in undamaged *brat^1^/+* discs (Figure S3A-E). When damage is induced, Wg is upregulated throughout the blastema by R0 (23) (Figure 3C). Reduction of *brat* expression resulted in significantly higher levels of Wg expression at R0 (Figure 3D and 3E) but not at R24 (Figure 3F). After ablation, Myc expression is elevated in the regenerating tissue (23) (Figure 3G and 3H). *brat^1^/+* discs showed significantly higher levels of Myc at R0, which were sustained through R24 (Figure 3I-K). Thus, being heterozygous mutant for *brat* caused an increase in the levels of both Wg and Myc early in regeneration. The elevated expression of these growth regulators likely explains the high proliferation seen in *brat^1^/+* discs at R0, and the larger wing pouch at R48.

**Figure 3.**
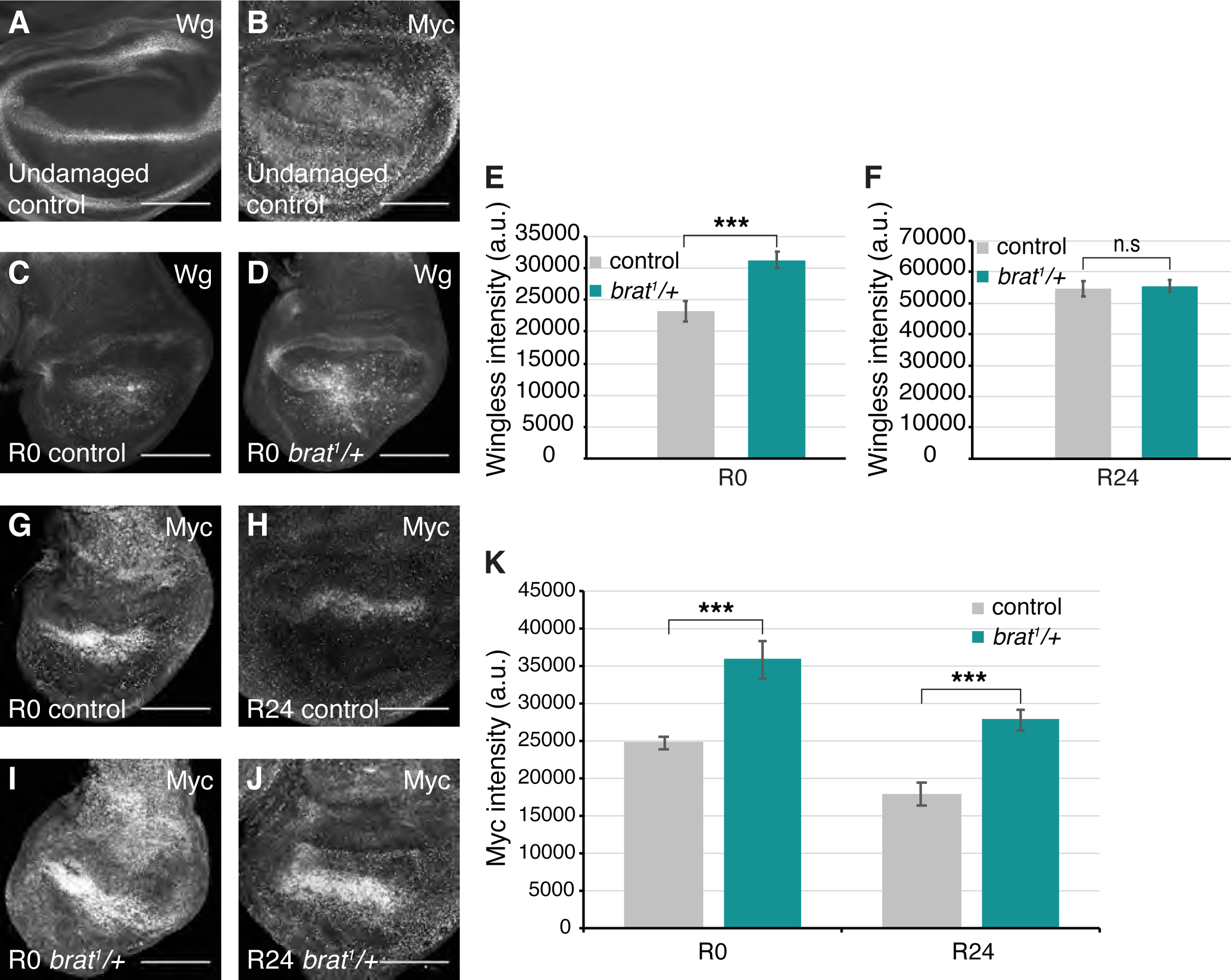
*brat^1^/+* animals experience elevated regeneration signaling. (A) Anti-Wg immunostaining in an undamaged control (*w^1118^*) disc. (B) Anti-Myc immunostaining in an undamaged control (*w^1118^*) disc. (C-D) Anti-Wg immunostaining in an R0 control (*w^1118^*) disc (C) and an R0 *brat^1^/+* disc (D). (E) Quantification of Wg fluorescence intensity in R0 control (*w^1118^*) (n = 13) and R0 *brat^1^/+* (n = 17) discs. *** p < 0.0006. (F) Quantification of Wg fluorescence intensity in R24 control (*w^1118^*) (n = 12) and R24 *brat^1^/+* (n = 11) discs. Area for fluorescence intensity measurement was defined by the Wg expression domain in the wing pouch. (G-J) Anti-Myc immunostaining in an R0 control (*w^1118^*) disc (G), an R24 control (*w^1118^*) disc (H), an R0 *brat^1^/+* disc (I) and an R24 *brat^1^/+* disc (J). (K) Quantification of Myc fluorescence intensity in R0 control (*w^1118^*) (n = 13), R0 *brat^1^/+* (n = 12), R24 control (*w^1118^*) (n = 13), and R24 *brat^1^/+* (n = 12) discs. Area for fluorescence intensity measurement was defined by the elevated Myc expression domain in the wing pouch. R0 *** p < 0.0003, R24 *** p < 0.0001. Error bars represent SEM. Student’s T-test used for statistical analyses. Scale bars are 100 μm.

### *brat* is required for margin cell-fate specification during regeneration

Reduction of *brat* during regeneration caused patterning defects specifically at the wing margin, resulting in the loss of vein at the margin and loss of sensory bristles (Figure 1G). Thus, *brat* is required for correct cell-fate specification at the DV boundary during regeneration. The wing imaginal disc is divided into the dorsal and the ventral compartments, with expression of the LIM-homeodomain protein Apterous (Ap) in dorsal cells. The juxtaposition of the dorsal and ventral cells forms the DV boundary, which develops into the adult wing margin (63) (Figure 4A). Notch (N) and Wg signaling at the DV boundary are crucial for the correct organization and cell-fate specification at the boundary (64). *cut (ct)* and *achaete (ac)* are margin-specific genes that are expressed downstream of N and Wg signaling. *ct* is required for the specification of the wing margin, and *ac* specifies the pro-neural sensory organ precursors (64,65) (Figure 4A).

**Figure 4.**
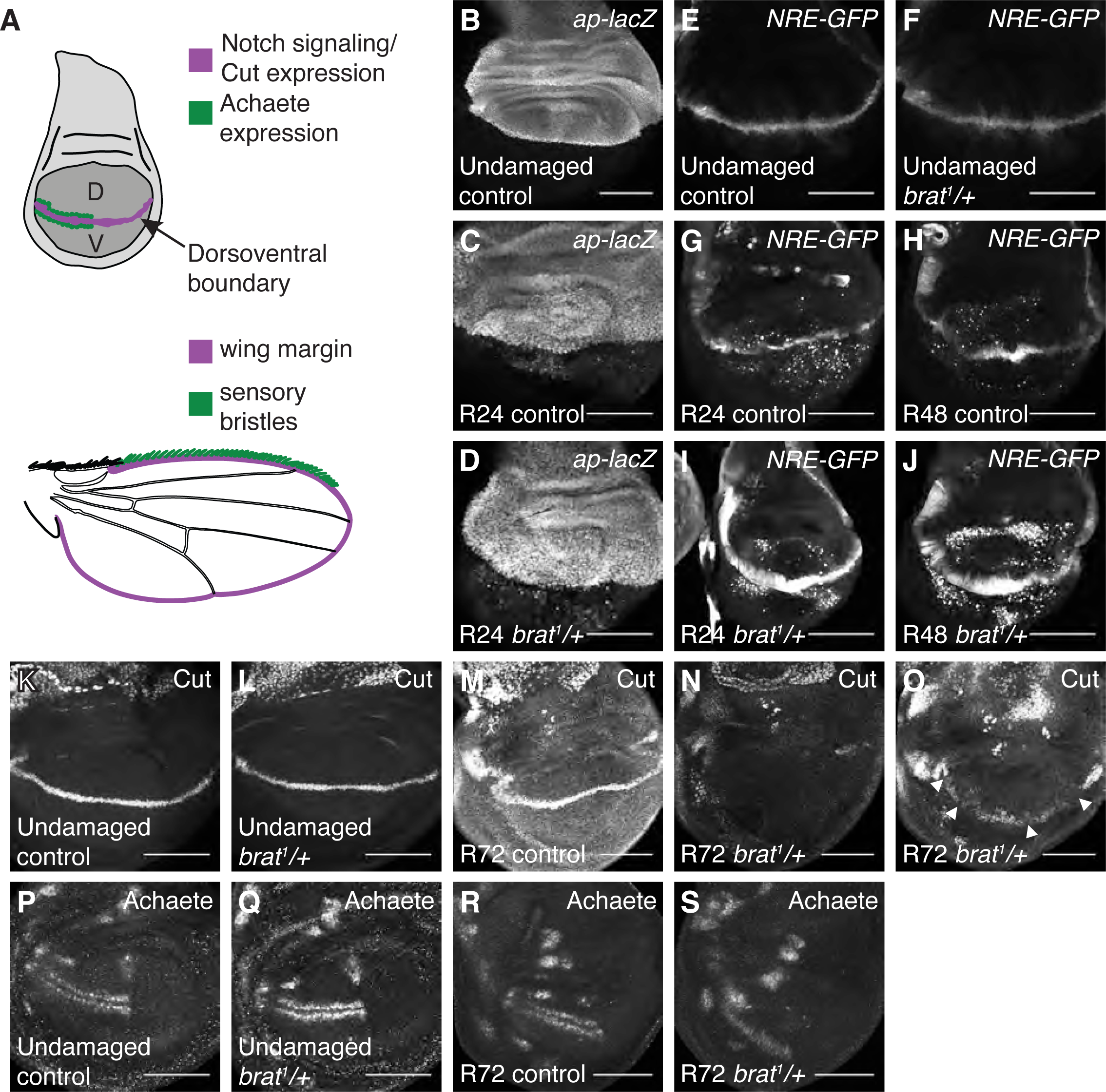
*Brat* regulates margin cell-fate specification. (A) Drawings of a wing imaginal disc and an adult wing. D = dorsal and V = ventral compartments of the wing disc, with the dorsoventral boundary marked in purple. Notch signaling and Cut expression are present at the dorsoventral boundary, which forms the adult wing margin, also marked in purple. Achaete-expressing cells, marked in green, give rise to the sensory bristles at the anterior half of the margin in the adult wing, also marked in green. (B) *ap-lacZ* expression in an undamaged control disc from a third- instar *ap-lacZ/CyO* animal. (C-D) *ap-lacZ* expression in an R24 control (*w^1118^*) disc (C) and an R24 *brat^1^/+* disc (D). (E-F) *NRE-GFP* expression in an undamaged control (*w^1118^*) disc (E) and an undamaged *brat^1^/+* disc (F). (G-J) *NRE-GFP* expression in an R0 control (*w^1118^*) disc (G), an R24 control (*w^1118^*) disc (H), an R0 *brat^1^/+* disc (I) and an R24 *brat^1^/+* disc (J). (K-L) Anti-Ct immunostaining in an undamaged control (*w^1118^*) disc (K) and an undamaged *brat^1^/+* disc (L). (M-O) Anti-Ct immunostaining in an R72 control (*w^1118^*) disc (M) and an R72 *brat^1^/+* discs (N-O). Arrowheads point to loss of Ct expression in (O). (P-Q) Anti-Ac immunostaining in an undamaged control (*w^1118^*) disc (P) and an undamaged *brat^1^/+* disc (Q). (R-S) Anti-Ac immunostaining in an R72 control (*w^1118^*) disc (R) and an R72 *brat^1^/+* disc (S). Scale bars are 100 μm.

To investigate whether the errors in fate specification seen in *brat^1^/+* discs were due to a compromised compartment boundary, we examined the expression of Ap using the *ap-lacZ* reporter. *ap-lacZ* expression showed a clear DV boundary in the undamaged control discs (Figure 4B). The DV boundary remained intact after ablation in control and *brat^1^/+* discs (Figure 4C-D, Figure S4A and S4B). *ap-lacZ* expression was also seen in the debris found in the damaged wing imaginal disc, due to the perdurance of β-gal. Wg expression was restored to its normal DV expression by R48 in both control and *brat^1^/+* discs (Figure S4C and S4D). Therefore, the patterning defects were not caused by disruptions in the DV boundary or changes in Wg expression.

Next, we examined N signaling in *brat^1^/+* discs due to its critical role in specifying fates at the DV boundary. We used a N signaling reporter, which uses *Notch Response Elements* (*NREs*) that bind to the Notch co-receptor Suppressor of Hairless, to drive the expression of GFP (66). No difference was detected in the expression of the N reporter for undamaged control and *brat^1^/+* discs (Figure 4E and 4F). N signaling at the DV boundary was restored by R24 in controls and continued at R48 (Figure 4G and 4H).

Note that the reporter signal can also be seen in cellular debris in the regenerating discs due to the perdurance of GFP. Interestingly, *brat^1^/+* discs showed highly elevated levels of the N signaling reporter at both these time points (Figure 4I and 4J). This result is consistent with recent evidence demonstrating Brat’s ability to attenuate N nuclear transport in the brain (67). We wondered whether this elevated N signaling could also disrupt margin fates. However, overexpressing the N-intracellular domain in the wing pouch during the 24-hour ablation period (Figure S4E and S4F) resulted in adult wings that were patterned remarkably well, with significantly fewer wings showing any margin defects when compared to the control (Figure S4G). Thus, increased N activity during regeneration suppresses margin defects. Additionally, decreasing N signaling activity in *brat^1^/+* regenerating discs by using a mutation in the *anterior pharynx defective 1 (aph- 1)* gene was unable to rescue the *brat* heterozygous phenotype. *aph-1^D35^/+* discs showed significantly reduced N signaling during normal development (Figure S4H-J) and at R24 during regeneration (Figure S4K-M), but could not rescue the loss of margin phenotype in the *brat* mutant (Figure S4N). Thus, while Brat constrains N signaling during regeneration, the elevated N signaling in *brat^1^/+* mutants does not cause the margin cell-fate specification defects.

To determine whether the loss of margin cell fates was due to apoptosis of those cells later in regeneration, we immunostained regenerating discs for the cleaved caspase Dcp-1 as a marker of cell death, and Wg to mark the margin. We did not observe dying cells in the margin in either control or *brat^1^/+* regenerating discs (Figure S4 C-D).

### *brat* specifies margin fate by controlling the expression of Cut and Achaete

To understand how patterning was disrupted in *brat^1^/+* regenerating discs, we examined expression of margin cell-fate genes. Cut (Ct) expression was present along the DV boundary in both undamaged control and *brat^1^/+* discs (Figure 4K and 4L), consistent with our results showing that adult undamaged *brat^1^/+* wings do not have margin defects (Figure S1D). In control regenerating discs, Ct expression was detected at the DV boundary at R72, which is when regeneration and repatterning are largely complete (Figure 4M). By contrast, Ct expression was either not observed in *brat^1^/+* discs or was still missing in segments of the DV boundary at R72 (Figure 4N and 4O). These results indicate a specific error in cell-fate specification, as the DV boundary was intact at R72 (Figure S4A and S4B). Undamaged control and *brat^1^/+* discs also showed appropriate Ac expression in two stripes of cells directly flanking the DV boundary in the anterior half of the disc (Figure 4P and 4Q). Ac expression was also detected in control regenerating discs at R72 (Figure 4R). While Ac-expressing cells appeared in *brat^1^/+* discs, they were not clearly separated across the DV boundary (Figure 4S). This finding is consistent with previous reports showing that Ct suppresses Ac at the margin, and mutations in *ct* lead to aberrant expression of Ac at the DV boundary, followed by degeneration of the wing margin through cell death (68,69).

### High Myc expression perturbs margin cell-fate specification during regeneration

Our results show that Brat both restricts regenerative growth and ensures correct cell-fate specification at the wing margin. Interestingly, JNK signaling in regenerating tissue can cause aberrant posterior-to-anterior cell-fate changes, which can be suppressed by a regeneration-specific protective factor, Taranis, to ensure correct patterning of the regenerating tissue (32). Therefore, we wondered whether unconstrained regenerative growth, or unconstrained expression of growth drivers, could also have deleterious side effects such as loss of margin cell fates. As Wg expression is normal during late regeneration and we have ruled out elevated N signaling as the causative factor for the cell-fate errors that occurred in *brat^1^/+* regenerating discs, we wondered whether high Myc levels caused by reduced Brat could cause the margin defects.

Brat overexpression can suppress Myc in wing imaginal disc cells (54), and in undamaged wing discs Brat protein levels were elevated at the DV boundary where Myc was reduced (Figure 5A and 5A’), suggesting that Brat may regulate Myc at the DV boundary. Furthermore, our results showed that regenerating *brat/+* mutant discs experienced elevated Myc levels compared to controls (Figure 3G-K). Previous studies demonstrated that Brat regulates Myc at the post-transcriptional level (54). To confirm that Brat is also regulating *myc* post-transcriptionally during regeneration, we measured *myc* transcript levels through qPCR. Regenerating discs showed significantly increased transcription of *myc* compared to undamaged controls. However, there was no significant difference in *myc* transcript levels between regenerating control and *brat^1^/+* discs at R0 and R24 (Figure 5B and 5C), indicating that Brat’s regulation of Myc must be post-transcriptional.

**Figure 5.**
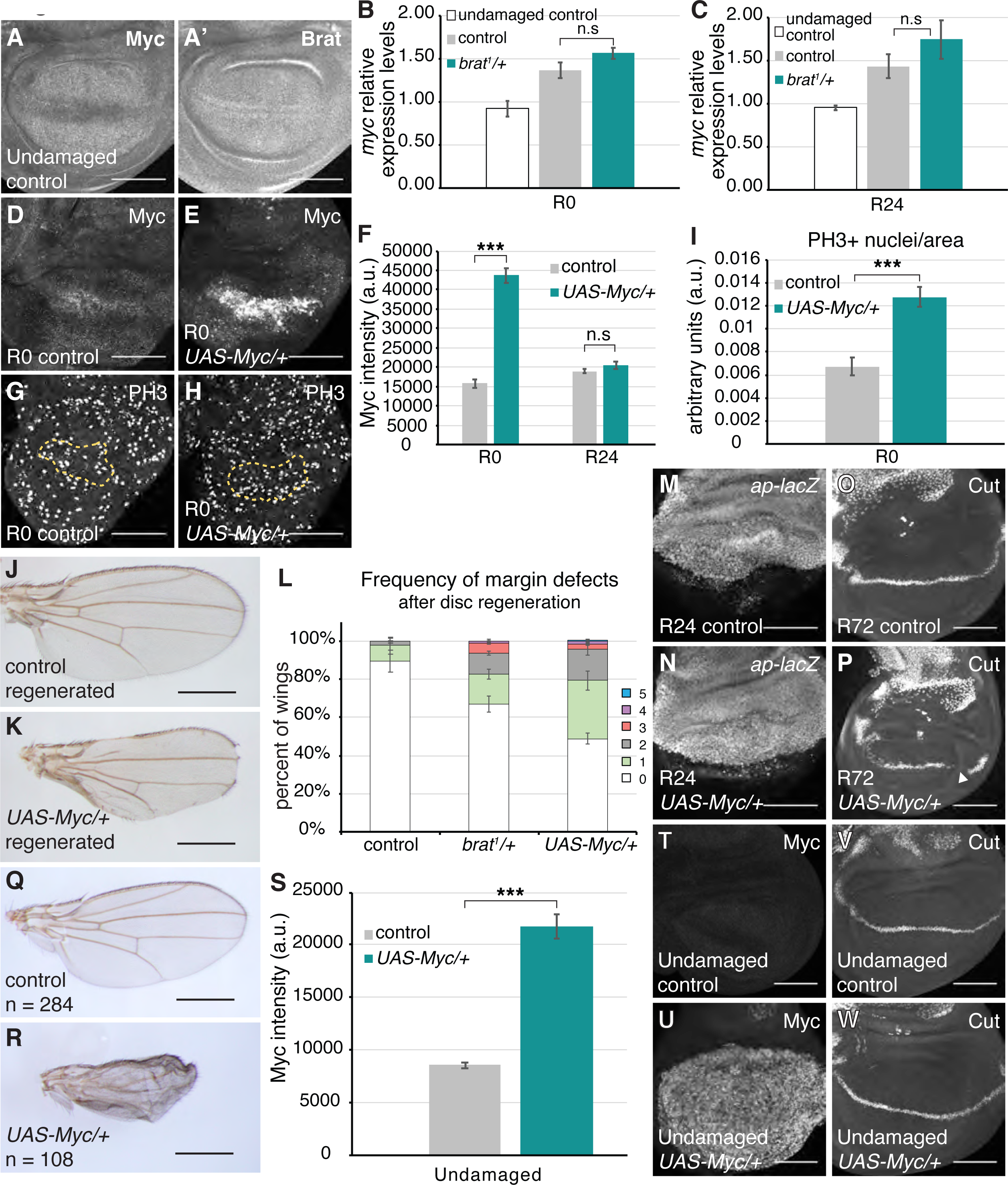
High Myc expression causes margin defects. (A-A’) Anti-Myc and Anti-Brat co-immunostaining in an undamaged control disc. *rnGAL4, GAL80^ts^/attP2* animals were shifted to 30°C on day 7 AEL and dissected 24 hours later. (B-C) Relative expression levels of *myc* for undamaged control, regenerating control (*w^1118^*), and regenerating *brat^1^/+* discs at R0 (B) and R24 (C). P values for comparison between regenerating control and *brat^1^/+* discs: p > 0.1 at R0, and p > 0.3 at R24. (D-E) Anti-Myc immunostaining in an R0 control (*w^1118^*) disc (D) and an R0 *UAS-Myc/+* disc (E). (F) Quantification of Myc fluorescence intensity in R0 control (*w^1118^*) (n = 13), R0 *UAS-Myc/+* (n = 12), R24 control (*w^1118^*) (n = 13), and R24 *UAS-Myc/+* (n = 12) discs. Area for fluorescence intensity measurement was defined by the elevated Myc expression domain in the wing pouch. *** p = 1.2E-11. (G-H) Anti-PH3 immunostaining in an R0 control (*w^1118^*) disc (G), and an R0 *UAS-Myc/+* disc (H). The yellow dashed lines outline the Nubbin-expressing wing pouch. (I) PH3-positive nuclei were counted within the regenerating wing pouch as marked by Anti-Nubbin co- immunostaining. Quantification of PH3-positive nuclei in the Nubbin area for R0 control (*w^1118^*) (n = 15) and *UAS-Myc/+* (n = 15) discs. *** p < 0.00002. (J) Adult control (*w^1118^*) wing after disc regeneration. (K) Adult *UAS-Myc/+* wing after disc regeneration. (L) Frequency of margin defects, as quantified in Figure 1H, seen in adult wings after disc regeneration for control (*w^1118^*) (n = 134), *brat^1^/+* (n = 193) and *UAS-Myc/+* (n = 200) wings, from three independent experiments. (M-N) *ap-lacZ* expression in an R24 control (*w^1118^*) disc (M) and an R24 *UAS-Myc/+* disc (N). (O-P) Anti-Ct immunostaining in an R72 control (*w^1118^*) disc (O) and an R72 *UAS-Myc/+* disc (P). (Q) Adult undamaged control (*+; rnGAL4,GAL80ts/+*) wing from animals shifted to 30°C on day 7 AEL and maintained at 30°C until eclosion. (R) Adult wing from discs continuously overexpressing Myc from day 7 AEL onwards (*UAS-Myc/+; rnGAL4, GAL80ts/+)*. (S) Quantification of Myc fluorescence intensity in undamaged discs dissected 31 hours after animals were shifted to 30°C on day 7 AEL. Control (*+; rnGAL4,GAL80ts/+*) (n = 14) and *UAS-Myc (UAS-Myc/+; rnGAL4, GAL80ts/+)* (n = 14). Area for fluorescence intensity measurement was defined by the Myc expression domain in the wing pouch. *** p = 1.2E-11. (T-U) Anti-Myc immunostaining in an undamaged control disc (T) and an undamaged *UAS-Myc/+* disc (U). (V-W) Anti-Ct immunostaining in an undamaged control disc (V) and an undamaged *UAS-Myc/+* disc (W). Error bars represent SEM. Student’s T-test used for statistical analyses. Scale bars are 100 μm. Scale bars for adult wings are 0.5 mm.

To test whether the elevated Myc protein levels could cause margin defects during regeneration and phenocopy the *brat* mutation, we overexpressed Myc in the wing pouch during the 24-hour ablation period. Myc was highly upregulated at R0 (Figure 5D- F), but Myc levels had returned to normal by R24 (Figure 5F). Overexpression of Myc also resulted in a significantly higher number of proliferating nuclei in the regenerating tissue at R0, similar to *brat^1^/+* discs (Figure 5G-I). Remarkably, we observed that adult wings resulting from Myc-overexpressing regenerating discs also showed margin defects similar to the *brat^1^/+* wings (Figure 5J and 5K). Moreover, the frequency of margin defects in the adult wings resulting from Myc-overexpressing regenerating discs was even higher than in adult wings resulting from *brat^1^/+* regenerating discs (Figure 5L). This difference in severity of phenotype correlates with levels of Myc at R0; in *brat^1^/+* regenerating discs Myc levels are higher than the control by 1.4 and 1.5-fold at R0 and R24 respectively (Figure 3K), and in Myc-overexpressing regenerating discs, Myc levels are higher than control by 2.75 fold at R0 but become similar to controls by R24 (Figure 5F). Thus, elevated levels of Myc alone can cause errors in margin cell-fate specification. Similar to *brat^1^/+* discs, *ap-lacZ* expression showed that the compartment boundary was not compromised in Myc-overexpressing regenerating discs (Figure 5M and 5N). Likewise, Ct expression was missing in segments at the DV boundary as in the *brat^1^/+* discs (Figure 5O and 5P).

Overexpressing Myc for a 24-hour window during normal development resulted in 3 adult wings out of 730 that showed any margin defects (Figure S5A). Even in these wings, only one segment of the margin was affected. To rule out the possibility that transient overexpression of Myc during development may not be sufficient to perturb patterning, we overexpressed Myc continuously after the animals entered the third instar larval stage. Continuous Myc overexpression proved to be lethal for many animals. While the flies that survived had significantly smaller wings, the margins showed no defect (Figure 5Q and 5R). 31 hours of Myc overexpression during normal development showed significantly higher Myc protein levels (Figure 5S-U) but did not interfere with Ct expression (Figure 5V and 5W). These data indicate that high Myc levels do not cause cell-fate specification errors during normal development, and the extensive loss of wing margin induced by high Myc expression is a regeneration-specific phenotype.

### Loss of Myc expression also perturbs margin cell-fate specification during regeneration

We hypothesized that if the *brat* phenotype was due to elevated Myc levels, we would be able to rescue the phenotype by reducing Myc levels in the *brat* mutant. For this purpose, we used *dm^4^*, which is a null allele of Myc (70). Surprisingly, we observed that the *dm^4^/+* mutants alone showed margin defects in the regenerated wings at a frequency similar to *brat^1^/+*, even though the *dm^4^/+; brat^1^/+* double mutant showed slightly reduced frequency of margin defects (Figure S5B). To confirm that Myc levels were reduced in the *dm^4^/+* mutants, we quantified Myc protein through immunostaining. We observed that there was no significant difference in Myc expression levels between the *dm^4^/+* mutant and control, both during development and regeneration (Figure S5C and S5D). Indeed, Myc levels were trending higher in the *dm^4^/+* discs during regeneration, perhaps accounting for the margin defects. The failure of the *dm^4^*mutation to reduce Myc levels could be due to compensatory expression of the functional copy of the Myc locus. We next tried reducing Myc levels though RNAi.

Despite the RNAi expression being transient in our system, and only occurring in cells that survive ablation, RNAi-mediated persistent knockdown has worked for multiple genes, likely due to the shadow RNAi effect (71). Two RNAi lines could significantly reduce Myc levels during normal development when expressed during early third instar (Figure S5E). However, when Myc RNAi was expressed during the 24-hour ablation period, Myc levels were not reduced at either R0 or R24, with one Myc RNAi line showing significantly higher levels of Myc compared to the control (Figure S5F).

To reduce Myc levels without the possibility of compensatory expression from an endogenous allele, we turned to the viable hypomorphic allele *Myc^P0^*, which is reported to eliminate Myc mRNA in the wing pouch (72). Indeed, we found that male larvae hemizygous for *Myc^P0^* did not show the typical Myc expression in third-instar wing discs, and lacked upregulation of Myc during regeneration (Fig S5 G-H). The *Myc^P0^*males developed more slowly than control larvae, so we adjusted the timing of induction of ablation later by 4 hours to coincide with the early third instar larval phase. Surprisingly, we found that *Myc^P0^* males that had undergone ablation and regeneration in the wing discs also had adult wings that were missing substantial sections of their margins, as did *Myc^P0^*; *brat^1^/+* double mutants (Fig S5I). Indeed, immunostaining *Myc^P0^* regenerating discs showed gaps in *cut* expression, indicating disruption of margin fate (Fig S5J-K).

Therefore, reduction of Myc as well as elevation of Myc disrupts margin cell fate during regeneration, underscoring the importance of tightly regulating expression of this pro-regeneration, pro-growth factor. However, this phenotype means that rescuing the *brat^1^/+* phenotype by reducing Myc is not possible, given that reducing Myc alone disrupts the wing margin, and we cannot rule out the possibility that other Brat targets in addition to Myc contribute to the patterning defects in *brat^1^/+* regenerated wings.

Importantly, animals that overexpressed Myc in the wing pouch during ablation did not undergo a regeneration-induced pupariation delay (Figure S5L), confirming that Brat regulates the entry into metamorphosis independently of its regulation of Myc.

Therefore, not all effects of reduced Brat are mediated through Myc.

### Enhanced proliferation does not disrupt margin cell-fate specification during regeneration

Myc is an important driver of regenerative growth, and yet, we found that cell-fate specification during regeneration can be negatively affected if Myc levels are not constrained. To test whether the aberrant patterning was a specific result of high Myc levels or whether increases in growth and proliferation could, in general, cause margin defects, we also increased proliferation during regeneration by overexpressing both *cyclinE (cycE)* and *string (stg)*.

Overexpressing the cell cycle genes in the wing imaginal disc during the 24-hour ablation period caused the resulting adult wings to be much larger than controls that had also undergone damage and regeneration (Figure 6A), though not larger than a normal wing. No significant regeneration-induced pupariation delay was seen, making the enhanced regeneration even more remarkable (Figure 6B). Intriguingly, we did not observe many margin defects for wings that had experienced *cycE* and *stg* overexpression during regeneration (Figure 6C-E). Furthermore, we have not observed this loss of margin phenotype with overexpression of other pro-regeneration genes such as *dilp8* (data not shown). Thus, pattern disruption during regeneration appears to be specifically associated with Myc overexpression and does not appear to be caused by increased proliferation alone.

**Figure 6.**
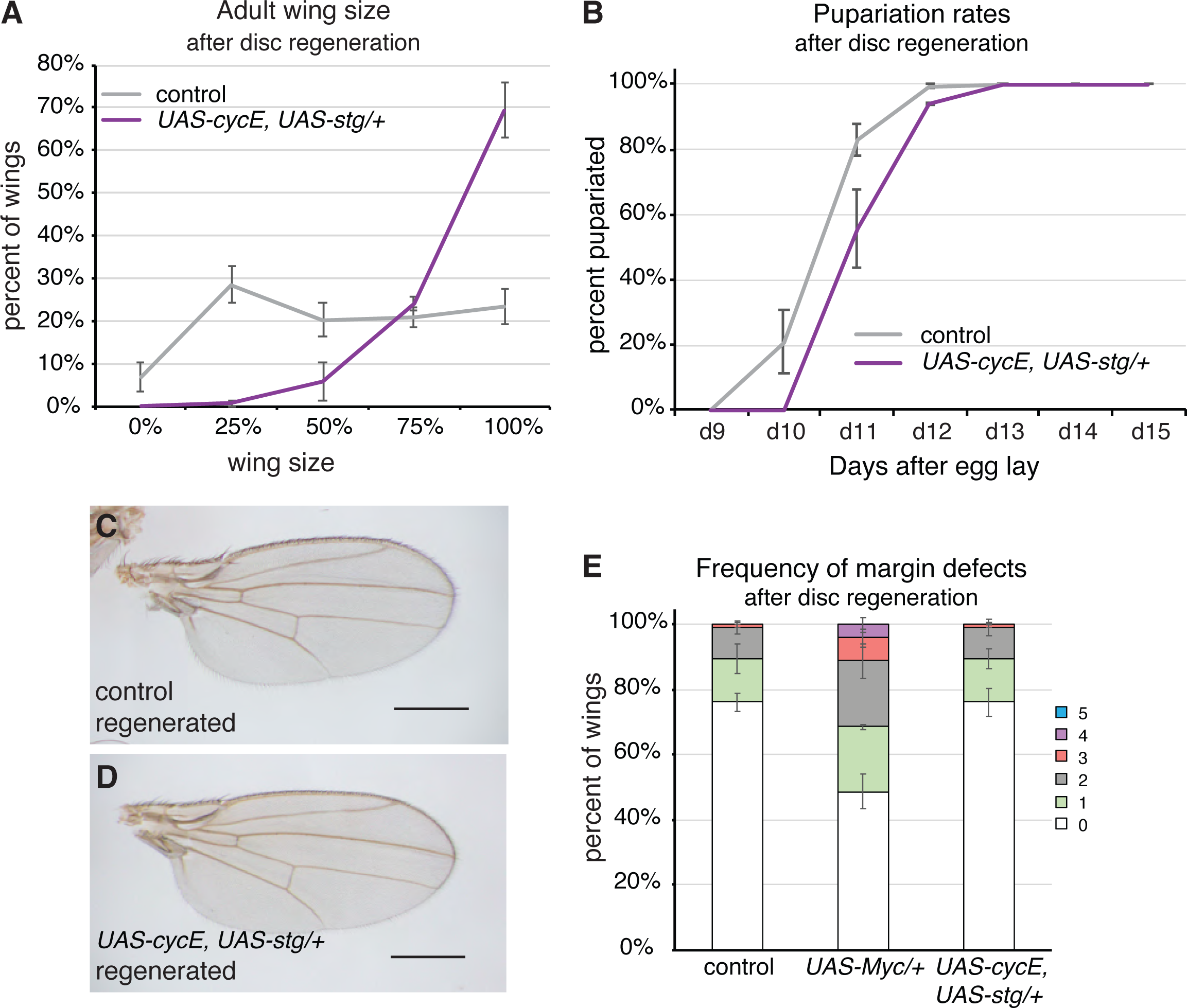
Overexpression of cell cycle genes does not cause patterning defects. (A) Adult wing sizes observed after disc regeneration for control (*w^1118^*) (n = 280) and *UAS-cycE, UAS-stg/+* (n = 194) wings, from three independent experiments. (B) Pupariation rates after disc regeneration for control (*w^1118^*) (n = 174) and *UAS-cycE, UAS-stg/+* (n = 115) wings, from three independent experiments. (C) Adult control (*w^1118^*) wing after disc regeneration. (D) Adult *UAS-cycE, UAS-stg/+* wing after disc regeneration. (E) Frequency of margin defects seen in adult wings after disc regeneration for control (*w^1118^*) (n = 118), *UAS-Myc/+* (n = 146), and *UAS-cycE, UAS- stg/+* (n = 188) wings, from three independent experiments. Error bars represent SEM. Scale bars for adult wings are 0.5 mm.

### Loss of cell-fate specification may be due to misregulation of Myc target genes

Given that driving growth by overexpressing Cyclin E and String does not cause loss of wing margin cell fates in regenerating tissue, this phenotype might be caused by misregulation of one or more targets of the Myc transcription factor. To identify genes that may regulate margin cell fate downstream of Myc, we screened through genes reported in the literature as regulating *ct* expression, and identified one that was misregulated in *brat^1^/+* and *UAS-Myc* regenerating discs. We have previously identified the gene *Chronologically inappropriate morphogenesis (chinmo)* as a novel regulator of regeneration (50). Chinmo is a transcription factor that regulates the balance between a proliferative self-renewal state and a differentiated state in stem cells (73,74). Recent work has shown that *chinmo* also maintains wing epithelial cells in an unspecified state during development by inhibiting *ct* expression, and enhances regenerative potential (75). Therefore, we included *chinmo* in our targeted screen for genes that are misregulated in the *brat^1^/+* or *UAS-Myc* regenerating discs and that contribute to the disruption of cell fate.

Several lines of evidence suggest that *chinmo* can be a transcriptional target of Myc. First, the model organism Encyclopedia of Regulatory Networks (modERN) data show Myc binding near the *chinmo* promoter (76). Second, ChIP-seq experiments in *Drosophila* embryonic Kc cells have shown that Myc does indeed bind to the *chinmo* locus (77). Finally, *chinmo* levels are upregulated in response to elevated Myc expression in wing imaginal discs (78). To determine whether *chinmo* expression was altered in *brat^1^/+* wing discs, we immunostained for Chinmo protein. Chinmo levels were not significantly different in undamaged control and *brat^1^/+* discs (Figure 7A, 7B and 7E). However, Chinmo levels were significantly higher in *brat^1^/+* regenerating discs compared to control regenerating discs at R24 (Figure 7C, 7D and 7F). To confirm regulation of *chinmo* downstream of Myc, we examined Chinmo levels in regenerating discs over-expressing Myc. Chinmo levels were elevated in Myc-overexpressing discs at R0, when Myc overexpression was the highest (Figure 5F and Figure 7G-I).

**Figure 7.**
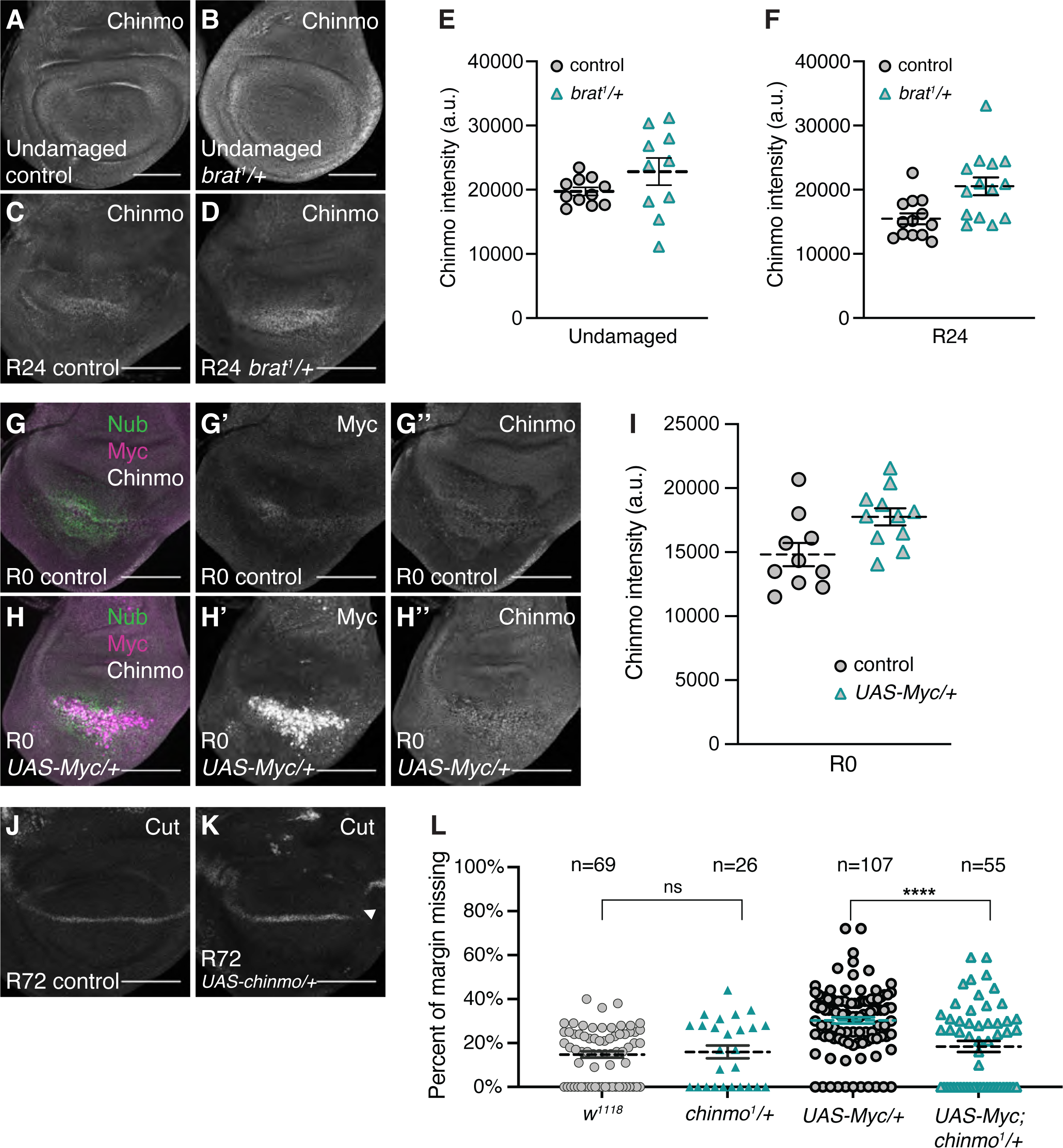
Chinmo levels are elevated in *brat^1^/+* and Myc-overexpressing regenerating discs. (A-B) Anti-Chinmo immunostaining in an undamaged control (*w^1118^*) disc (A) and an undamaged *brat^1^/+* disc (B). (C-D) Anti-Chinmo immunostaining in an R24 control (*w^1118^*) disc (C) and an R24 *brat^1^/+* disc (D). (E) Quantification of Chinmo fluorescence intensity in undamaged control (*w^1118^*) (n = 11) and undamaged *brat^1^/+* (n = 10) discs. p=0.16 (F) Quantification of Chinmo fluorescence intensity in R24 control (*w^1118^*) (n = 13) and R24 *brat^1^/+* (n = 14) discs. p < 0.006. Area for fluorescence intensity measurement was defined by the elevated Chinmo expression domain in the wing pouch. (G) Merge of anti-Nubbin, anti-Myc and anti-Chinmo immunostaining in an R0 control (*w^1118^*) disc. (G’-G’’) Same disc as (G) showing anti-Myc and anti-Chinmo immunostaining, respectively. (H) Merge of anti-Nubbin, anti-Myc and anti-Chinmo immunostaining in an R0 *brat^1^/+* disc. (H’-H’’) Same disc as (H) showing anti-Myc and anti-Chinmo immunostaining, respectively. (I) Quantification of Chinmo fluorescence intensity in R0 control (*w^1118^*) (n = 10) and R0 *UAS-Myc/+* (n = 11) discs. p < 0.02. Area for fluorescence intensity measurement was defined by the elevated Myc expression domain in the wing pouch. (J-K) Anti-Cut immunostaining in control (J) and UAS-chinmo (K) R72 discs. Arrowhead shows missing Cut expression in K. (L) Margin tissue lost as a percentage of total wing perimeter for control (*w^1118^*) (n = 69), *chinmo^1^/+ (n=26), UAS-Myc/+* (n = 107), and *UAS-Myc/+; chinmo^1^/+* (n=55) wings. ****p = 9.11E-05. Error bars represent SEM. Student’s T-test used for statistical analyses. Scale bars are 100 μm.

However, Chinmo levels were restored to control levels by R24 in Myc-overexpressing discs, consistent with the return of Myc levels to normal at this time point (Figure 5F and Figure S6A-C). Interestingly, Myc and Chinmo expression almost perfectly co-localized, consistent with the hypothesis that Myc regulates Chinmo expression (Figure S6A-B’’’). Additionally, we observed a high correlation between Myc and Chinmo expression levels in individual discs (Figure S6D and S6E). Undamaged discs overexpressing Myc did not show elevated Chinmo levels (Figure S6F-H), possibly explaining why Myc overexpression during normal development did not cause margin defects.

While *chinmo* mRNA can be a direct Brat target (39), *chinmo* is regulated at the level of transcription in the wing imaginal disc (75). To confirm the absence of regulation of *chinmo* mRNA by Brat during regeneration, we assessed expression of mCherry from a transgene in which the *chinmo* UTRs were fused to the *mCherry* coding region to serve as a reporter for *chinmo* mRNA regulation (*UAS-mCherry-chinmoUTR*) (79). We did not detect any change in mCherry levels in *brat^1^/+* regenerating discs compared to control regenerating discs, indicating that reduction of Brat does not impact the *chinmo* mRNA reporter and Brat does not likely regulate *chinmo* mRNA in regenerating discs (Fig S6I).

Thus, the loss of *ct* expression and loss of margin cell fates in *brat/+* regenerating discs and *UAS-Myc* regenerating discs may be due, at least in part, to upregulation of Myc targets, including *chinmo*. Importantly, other unidentified Myc targets likely also contribute to *ct* misregulation in regenerating discs with reduced Brat or overexpressed Myc. To test whether expression of *chinmo* in the regeneration blastema is sufficient to disrupt the margin, we expressed *UAS*-*chinmo* alongside the ablation system. Doing so led to a moderate level of gaps in cut expression in R72 regenerating discs (4/14 compared to 0/12 for controls) (Figure 7J, K). To test whether the loss of margin in regenerating wing discs overexpressing Myc is due at least in part to expression of *chinmo*, we asked whether the phenotype could be ameliorated by a *chinmo* mutation. Indeed, regenerating wing discs overexpressing Myc but heterozygous for *chinmo^1^* had significantly less margin loss compared to those just overexpressing Myc (Figure 7L).

Thus, we have identified at least one Myc target that contributes to disruption of cell fate in regenerating tissue with Myc expression elevated beyond what is normally found in the regeneration blastema.

Based on our findings, we propose a model in which pro-growth factors are important for coordinating regenerative growth, but can lead to deleterious side effects by perturbing cell-fate gene expression and patterning. Brat prevents a prolonged proliferative and unspecified state in regenerating wing discs by constraining Wg, Ilp8, Myc, and Myc targets, to enable cessation of growth, induction of cell-fate specification, and entry into metamorphosis (Figure 8).

**Figure 8.**
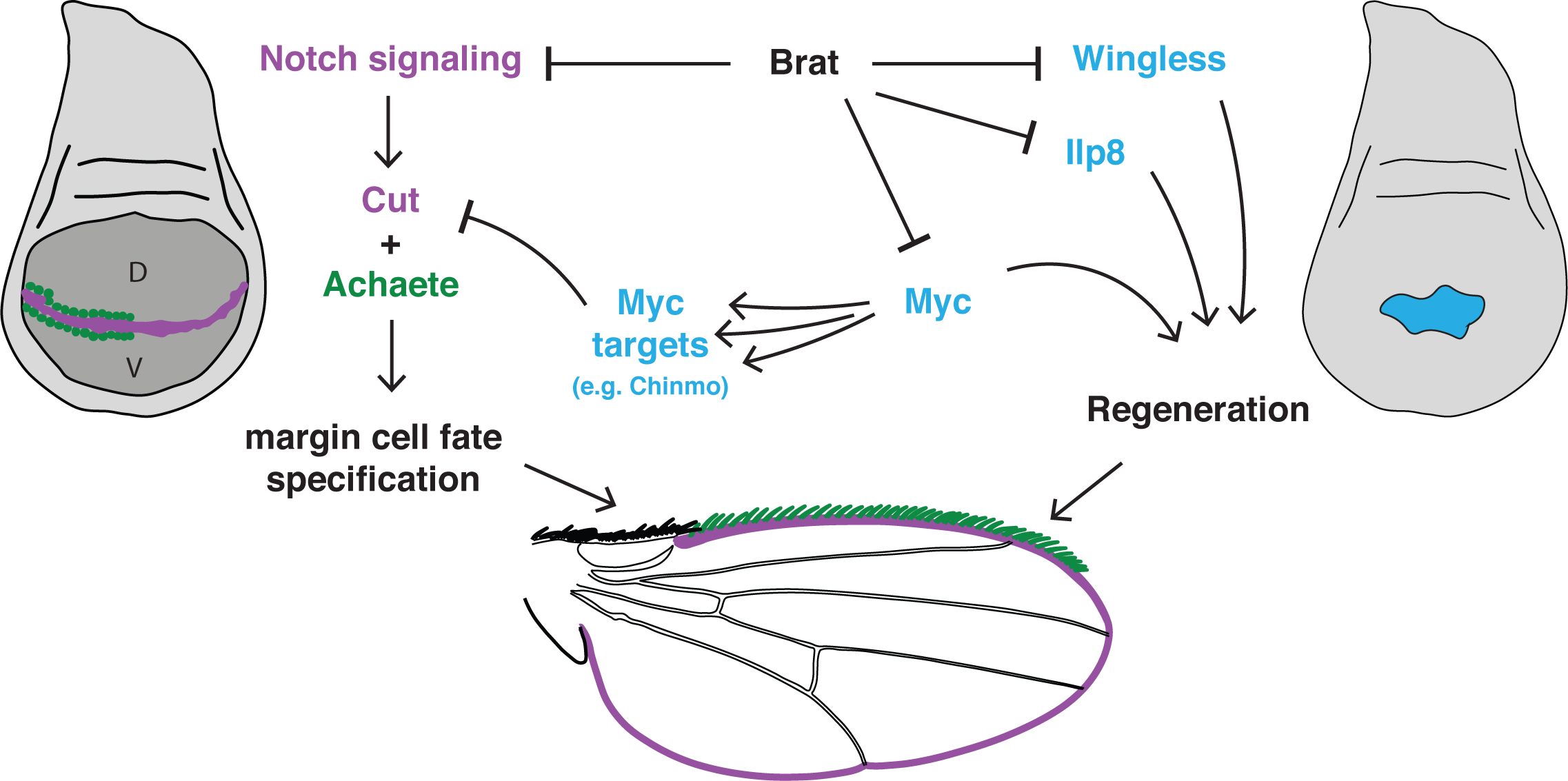
Brat restricts pro-regeneration factors and ensures correct margin cell- fate specification. Model describing the network of Brat targets in the regenerating wing imaginal disc. Importantly, Brat restricts Myc levels, limiting expression of Myc’s targets, including Chinmo, to allow correct margin cell-fate specification.

## Discussion

Here we have shown that Brat acts as a protective factor during regeneration, in part by constraining levels of the transcription factor Myc, which promotes growth and proliferation but also inhibits cell-fate specification. If Brat is unable to perform its protective function during regeneration, Myc levels increase unchecked, resulting in misregulation of its targets, causing loss of proper cell fates at the wing margin.

Myc is broadly used across organisms to promote proliferation and prevent differentiation (80,81). It is is required for efficient regeneration, and increased levels can enhance regeneration in both younger discs as well as mature discs that normally regenerate poorly (23,25). Nevertheless, we have found that while these abnormally high Myc levels can enhance regenerative growth, they can also perturb differentiation by misregulating target genes such as Chinmo. Thus, enhanced regeneration happens at the expense of correct cell-fate specification. Intriguingly, our attempts to rescue the *brat/+* phenotype by reducing Myc led to the discovery that loss of Myc expression in the regeneration blastema also leads to loss of margin cell fates. Many Myc targets are ribosomal protein genes, and, interestingly, adults transheterozygous for mutations in the ribosomal protein gene *RpL38* also have missing wing margins even without damage and regeneration of the wing disc (82). This finding provides precedent for disruption of margin fates when ribosomal protein expression is reduced, and suggests that the reasons for the lost margin in the *Myc^P0^* and *UAS-Myc* regenerating discs may be different. Importantly, expression of *ct* and possibly other cell-fate genes requires an appropriate amount of Myc, with too much or too little Myc disrupting cell fate. This phenomenon of requiring just the right amount of signaling or gene activity is not unprecedented, as we have previously shown that activation of JNK signaling after wing disc damage requires the right amount of reactive oxygen species (ROS) production; too much ROS and too little ROS both lead to inadequate JNK activity and poor regeneration (51).

Brat promotes differentiation in *Drosophila* larval neuroblasts and ovarian germline stem cells by asymmetrically segregating to one of the daughter cells where it post-transcriptionally inhibits Myc (41,42). This daughter cell is then able to differentiate while the other daughter cell remains a stem cell. In *brat* mutants, progeny of stem cells are unable to differentiate, resulting in an abnormal expansion of the stem-cell population, which can form tumors in the brain (40–43). Thus, Brat protects these tissues from overproliferation of stem cells. Importantly, wing imaginal disc regeneration is not stem- cell based, but in wing disc regeneration Brat also inhibits Myc to allow correct cell-fate specification. Based on these similarities in function, Brat likely acts as a protective factor across different biological contexts, including regeneration that does not employ stem cells.

We have previously shown that JNK signaling can induce posterior-to-anterior fate changes in regenerating wing discs, which can be prevented by the protective factor Taranis (32). In addition, we have shown that the chromatin-remodeling SWI/SNF BAP complex, defined by the complex-specific component Osa, similarly prevents aberrant posterior-to-anterior fate changes in regenerating wing discs (83). We have now identified a third protective factor, Brat, which is needed specifically for correct patterning of the regenerating wing margin. Interestingly, while elevated JNK signaling causes anterior markers to appear in the posterior wing compartment, it does not cause margin loss, indicating that posterior fate and margin fate are regulated in distinct ways (32). Protective factors such as Tara, Osa, and Brat are important for maintaining the balance between fate specification and regenerative potential, but they do so by using very different mechanisms. While the molecular function of Tara is unknown, genetic interactions in *Drosophila* coupled with the demonstrated functions of its vertebrate homologs suggest it may regulate gene expression at the level of transcription and chromatin (84–87). By contrast, Brat acts as a translational repressor by negatively regulating translation and stability of its target mRNAs (88,89). Tara is required to prevent fate changes induced by JNK signaling, which is necessary for wound repair and regeneration but is not required for the normal growth of the wing. By contrast, Myc is required for both development and regeneration of the wing disc, but is constrained by Brat only during regeneration.

An important open question in the field of regeneration is how patterning and cell-fate specification are regulated in regenerating tissue, and whether these mechanisms are different from the developmental program. Many studies have highlighted that regeneration must be distinct from development in some ways, because the damaged tissue already has complex patterning, and the wound-healing response causes strong activation of signaling pathways, some of which are not normally present in developing tissue (1,3,25,30–35). We are just beginning to identify regulators like Brat that are crucial for attenuating regenerative growth signaling and shielding the regenerating tissue from the harmful side effects of such signaling. Identification of these regulators highlights the fact that the regenerating tissue is distinct from normally developing tissue. Since regeneration signaling is complex and comprised of many signaling pathways, additional factors that play protective roles during regeneration likely exist. Identification of these factors will be important for the development of clinical therapies targeted at tissue repair, enabling these therapies to protect against the deleterious side effects of exogenous and unconstrained pro-growth signaling.

## Materials and Methods

### Ablation and Regeneration experiments

Ablation experiments were done as previously described (32). Briefly, cell death was induced by driving *UAS-reaper* under *rotund-GAL4*, with *GAL80^ts^* for temporal control. Animals were raised at 18°C for 7 days after egg lay (AEL) (early third instar) before they were shifted to a 30°C circulating water bath for 24 hours. Animals were brought back to 18°C to allow regeneration. Wing discs were dissected at different time points after the end of ablation, or the animals were allowed to grow to adulthood to observe the adult wing phenotype. Undamaged control wing discs were the same genotype as the experimental animals but kept at 18°C and dissected on day 9 after egg lay, which is mid-late third instar. For undamaged adult wings, the animals were kept at 18°C until after eclosion. Any other undamaged conditions used are mentioned specifically in the figure legends.

### Fly stocks

The following *Drosophila* stocks were used: *w^1118^* (wild type)(90), *w^1118^ ; rnGAL4, UAS-rpr, tubGAL80ts/TM6B,tubGAL80* (23), *brat^1^* (91)(FBst0003988), *brat^192^* and *brat^150^* (52)(a gift from Juergen Knoblich, Austrain Academy of Science), *brat^11^* (53)(a gift from Chen-Yu Lee, University of Michigan), *Df(2L)Exel8040* (92)(FBst0007847), *Df(2L)TE37C-7* (93)(FBst0006089), *rnGAL4, tubGAL80ts/TM6B* (23)*, P{Trip.HM05078}attP2* (called *bratRNAi* in the text)(FBst0028590), *P{CaryP}attP2* (called *attP2* control in the text)(FBst0036303), *{PZ}ap^rK568^*(94)(FBst0005374), *NRE-GFP* (66)(FBst0030727), *UAS-Nintra* (a gift from Gary Struhl, Columbia University), *aph-1^D35^* (95)(FBst0063242), *UAS-Myc* (72)(FBst0009674), *UAS-cycE, stg* (a gift from Laura Buttitta, University of Michigan), *dm^4^* (70), *P{GD1419}v2947* (called *MycRNAi#1* in the text)(VDRC ID# 2947) and *P{GD1419}v2948* (called *MycRNAi#2* in the text)(VDRC ID# 2948), *P{GD6000}v15293* (called control in the text) (VDRC ID# 15293), *Myc^P0^* (72) (FBal0100372), UAS-mCherry^chinmoUTRs^ (73).All fly stocks are available from the Bloomington Drosophila Stock Center unless stated otherwise.

### Pupariation timing

Pupariation experiments were performed in a similar manner to the ablation experiments. Starting at day 9, newly formed pupal cases were counted in each vial. Pupal cases were counted every 24 hours, up until day 15. Pupariation rates from three independent experiments were used to calculate the average plotted in the graphs.

### Immunohistochemistry

Immunostaining was carried out as previously described (23). Primary antibodies were rat anti-Brat (1:200) (37) (a gift from Robin Wharton, Ohio State University), mouse anti- Nubbin (1:500) (96) (a gift from Steve Cohen, University of Copenhagen), rabbit anti- Phospho-Histone H3 (1:500) (Millipore), mouse anti-Wingless (1:100) (The Developmental Studies Hybridoma Bank [DSHB]), rabbit anti-dMyc (1:500 or 1:250) (Santa Cruz Biotechnologies), mouse anti-βgal (1:100) (DSHB), mouse anti-Cut (1:100) (DSHB), mouse anti-Achaete (1:10)(DSHB), rat anti-Chinmo (1:500) (a gift from Nick Sokol, Indiana University), and anti-cleaved Dcp-1 (1:250)(Cell Signaling #9578). The Developmental Studies Hybridoma Bank (DSHB) was created by the NICHD of the NIH and is maintained at the University of Iowa, Department of Biology, Iowa City, IA 52242. Secondary antibodies were AlexaFluor probes (1:1000) (Life Technologies). DNA was marked using TO-PRO3 (1:500) (Life Technologies) or DAPI (1:5000 of 0.5 mg/mL stock) (Sigma). Discs were mounted in Vectashield mounting medium (Vector Laboratories).

Discs were imaged on a Zeiss LSM 510 or a Zeiss LSM 700 confocal microscope. Parameters for imaging were identical for quantified images. Images were processed using ZEN lite (Zeiss), ImageJ (NIH) and Photoshop (Adobe). Maximum intensity projections were created for the confocal images. Fluorescence intensity was measured within the wing pouch as marked by anti-Nubbin or by using the morphology of the undamaged wing disc. Myc and Chinmo intensities were measured by outlining the region expressing elevated Myc or Chinmo levels. *NRE-GFP* intensity was measured by outlining the GFP-expressing region at the DV boundary.

### Adult wing quantifications

Adult wings were mounted in Gary’s Magic Mount (Canada balsam [Sigma] dissolved in methyl salicylate [Sigma]). Images were taken with an Olympus SZX10 microscope with an Olympus DP21 camera using the CellSens Dimension software (Olympus).

All adult wings that were 75% or 100% the size of a normal wing were used to quantify the loss of the wing margin. The wing margin was divided into five segments defined by where the wing veins intersect the margin. Each wing was scored for the number of segments with missing margin to assess the extent of the patterning defect.

Percentages from the three independent experiments were used to calculate averages plotted in the graphs. The area of undamaged and regenerated wings was measured using ImageJ (NIH). ImageJ was also used to measure the percentage of linear length of margin lost for the entire perimeter of the wing. Graphs were plotted using Excel and Graphpad Prism 7.

### qPCR

For quantitative PCR (qPCR), 40-60 wing imaginal discs were collected in Schneider’s medium and stored at -80°C. RNA was extracted using the Qiagen RNeasy Mini Kit (#74104), and cDNA synthesis was performed using the Superscript III First Strand Synthesis kit (#11752-050). qPCR reactions using the Power SYBR Green MasterMix (ABI) were run on the ABI Step One Plus Real Time PCR System. The experiment consisted of 3 biological replicates. For each biological replicate there were three technical replicates. Gene expression was analyzed by the ΔΔC_t_ method and normalized to *Gapdh2* expression. The following primers were used: *Gapdh2* forward primer (GTGAAGCTGATCTCTTGGTACGAC), *Gapdh2* reverse primer (CCGCGCCCTAATCTTTAACTTTTAC) (97), *ilp8* primers used from Qiagen (QT00510552), *dmyc* forward primer (AACGATATGGTGGACGATGG), and *dmyc* reverse primer (CGGCAGATTGAAGTTATTGTAGC) (98). For qPCR experiments undamaged controls were *rnGAL4, tubGAL80ts/TM6B* females crossed to *w^1118^* males and shifted to 30°C for 24 hours at 7 days AEL. Discs were dissected either immediately or 24 hours after shifting the animals back to 18°C for R0 and R24 time points, respectively.

## Acknowledgements

The authors would like to thank Amanda Brock and Sumbul Khan for critical reading of the manuscript and helpful discussions. We also thank Juergen Knoblich, Cheng-Yu Lee, Gary Struhl, Robin Wharton, Nick Sokol, Laura Buttitta, and Cedric Maurange for reagents. Stocks obtained from the Bloomington Drosophila Stock Center (NIH P40OD018537) were used in this study. Transgenic fly stocks were obtained from the Vienna Drosophila Resource Center (VDRC, www.vdrc.at). Antibiodies were obtained from the Developmental Studies Hybridoma Bank, created by the NICHD of the NIH and maintained at The University of Iowa, Department of Biology, Iowa City, IA 52242.

## Author Contribution Section

Experiments were designed by SNFA and RKSB, and conducted and analyzed by SNFA and FT-YH. The manuscript was written by SNFA and RKSB.

## Data Availability

All relevant data are available at (insert web address for UIUC library repository when manuscript accepted) and upon request.

---

**Figure S1.**
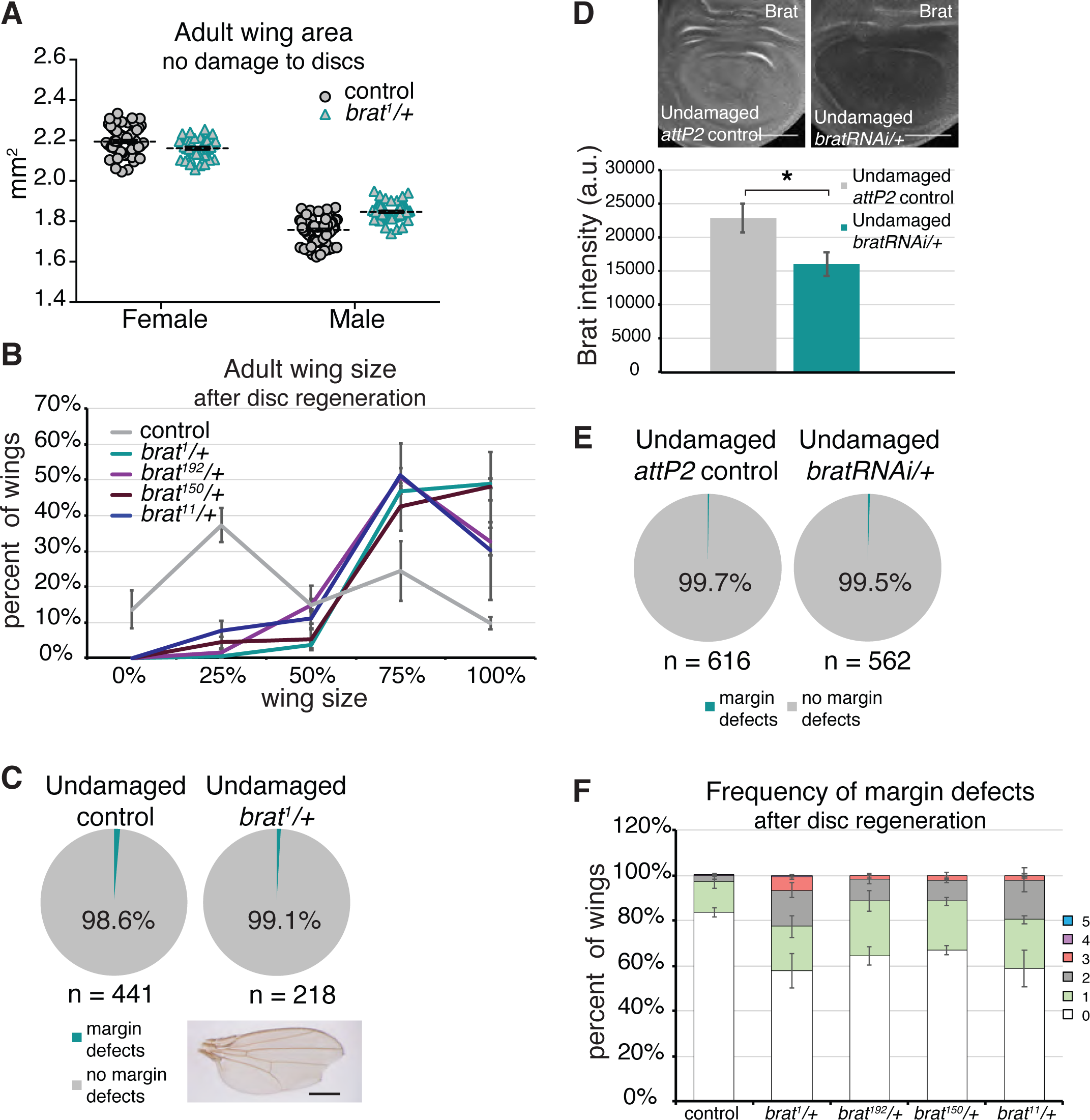
Reduction of *brat* expression does not cause enhanced growth or margin defects during normal development (Related to. **Figure 1**). (A) Adult wing area measured using ImageJ after mounting and imaging wings, for undamaged control (*w^1118^*) (n = 63 female and 70 male) and *brat^1^/+* (n = 38 female and 48 male) wings. *rnGAL4, GAL80^ts^/TM6B* females were crossed to *w^1118^* or *brat^1^/SM6- TM6B* males and taken through the protocol shown in Figure 1A. (B) Adult wing sizes after disc regeneration for control (*w^1118^*) (n = 599), *brat^1^/+* (n = 199), *brat^192^/+* (n = 237), *brat^150^/+* (n = 235) and *brat^11^/+* (n = 188) wings, from three independent experiments. (C) Margin defects detected in adult wings from undamaged control (*w^1118^*) and *brat^1^/+* discs. *rnGAL4, GAL80^ts^/TM6B* females were crossed to *w^1118^* or *brat^1^/SM6-TM6B* males and taken through the protocol shown in Figure 1A. Margin defects detected in the undamaged wings were never as severe as the ones seen in *brat^1^/+* wings after disc regeneration. A representative wing with margin defects is shown. (D) Anti-Brat immunostaining in undamaged control (*attP2*) and *bratRNAi/+* discs. *rnGAL4, GAL80^ts^/TM6B* females were crossed to *attP2* or *bratRNAi* males. Larvae were kept at 18°C and shifted to 30°C on day 7 AEL. Discs were dissected 24 hours after the shift to 30°C. Quantification of Brat fluorescence intensity in undamaged control *(attP2*) (n = 15) and *bratRNAi*/+ (n = 15) discs. Area for fluorescence intensity measurement was defined by wing pouch morphology and Anti-Myc co-immunostaining. * p = 0.02. (E) Margin defects detected in adult wings from undamaged control (*attP2*) and *bratRNAi* dics. *rnGAL4, GAL80^ts^/TM6B* females were crossed to *attP2* or *bratRNAi* males. Larvae were kept at 18°C and shifted to 30°C on day 7 AEL and kept there until eclosion. (F) Frequency of margin defects seen in adult wings after disc regeneration for control (*w^1118^*) (n = 240), *brat^1^/+* (n = 191), *brat^192^/+* (n = 196), *brat^150^/+* (n = 213) and *brat^11^/+* (n = 152) wings. Wings in (G) are from the same experiments as (B). Error bars represent SEM. Student’s T-test used for statistical analyses. Scale bars are 100 μm. Scale bars for adult wings are 0.5 mm.

**Figure S2.**
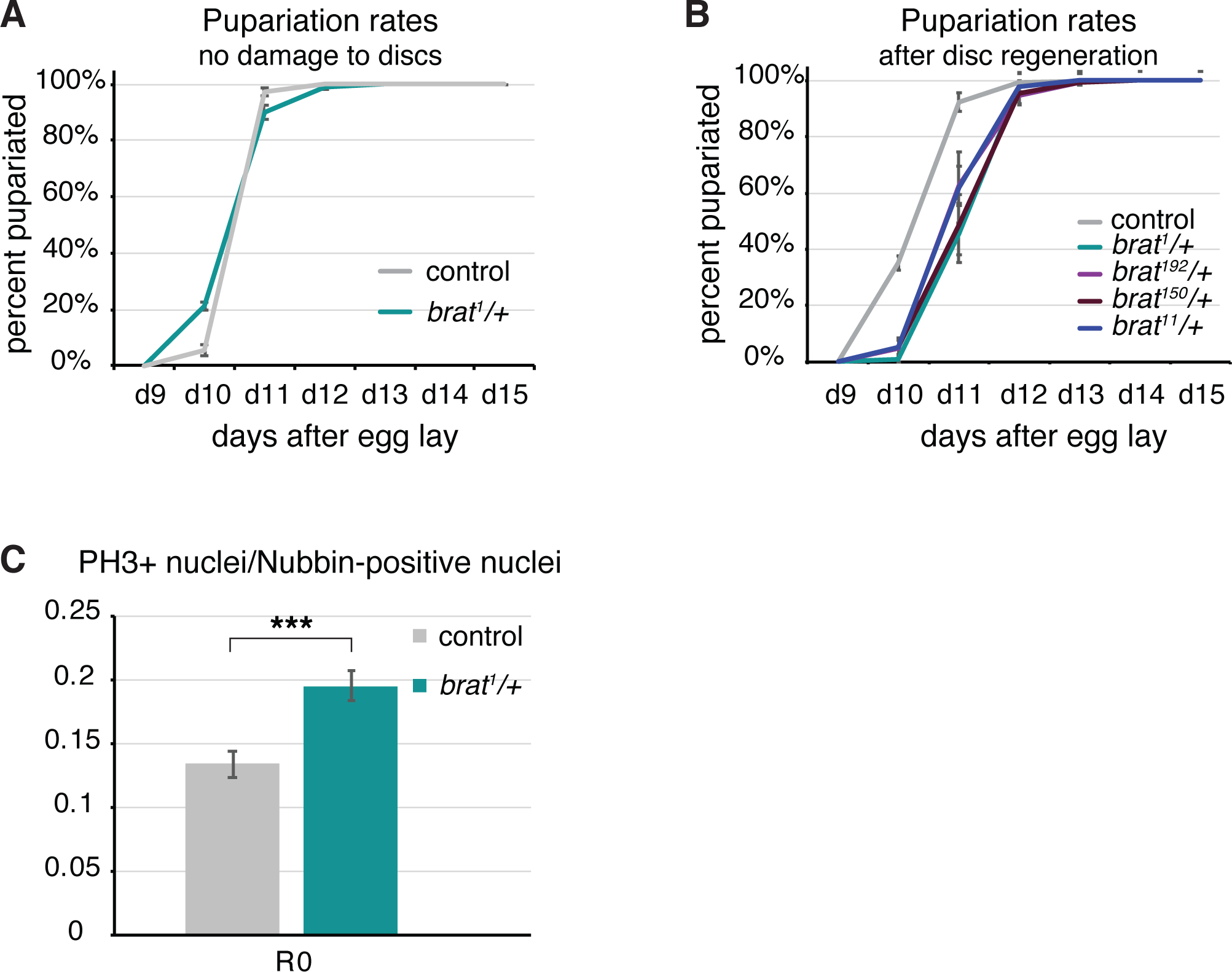
Reduction of *brat* expression delays pupariation in a regeneration- specific manner (Related to. **Figure 2**). (A) Pupariation rates in undamaged control (*w^1118^*) (n = 221) and *brat^1^/+* (n = 110) animals, from three independent experiments. (B) Pupariation rates after disc regeneration for control (*w^1118^*) (n = 384), *brat^1^/+* (n = 107), *brat^192^/+* (n = 131), *brat^150^/+* (n = 114) and *brat^11^/+* (n = 113) animals. Pupariation rates are from the same experiments as in Figure 2A. (C) PH3-positive nuclei were counted within the regenerating tissue as marked by Anti-Nubbin co-immunostaining. Total number of nuclei were also counted in the Nubbin expressing region. Ratio of PH3-positive nuclei and total Nubbin-positive nuclei for control (*w^1118^*) and *brat^1^/+* discs at R0 (n = 16 and 18). *** p < 0.0005. Error bars represent SEM. Student’s T-test used for statistical analyses. Error bars represent SEM.

**Figure S3.**
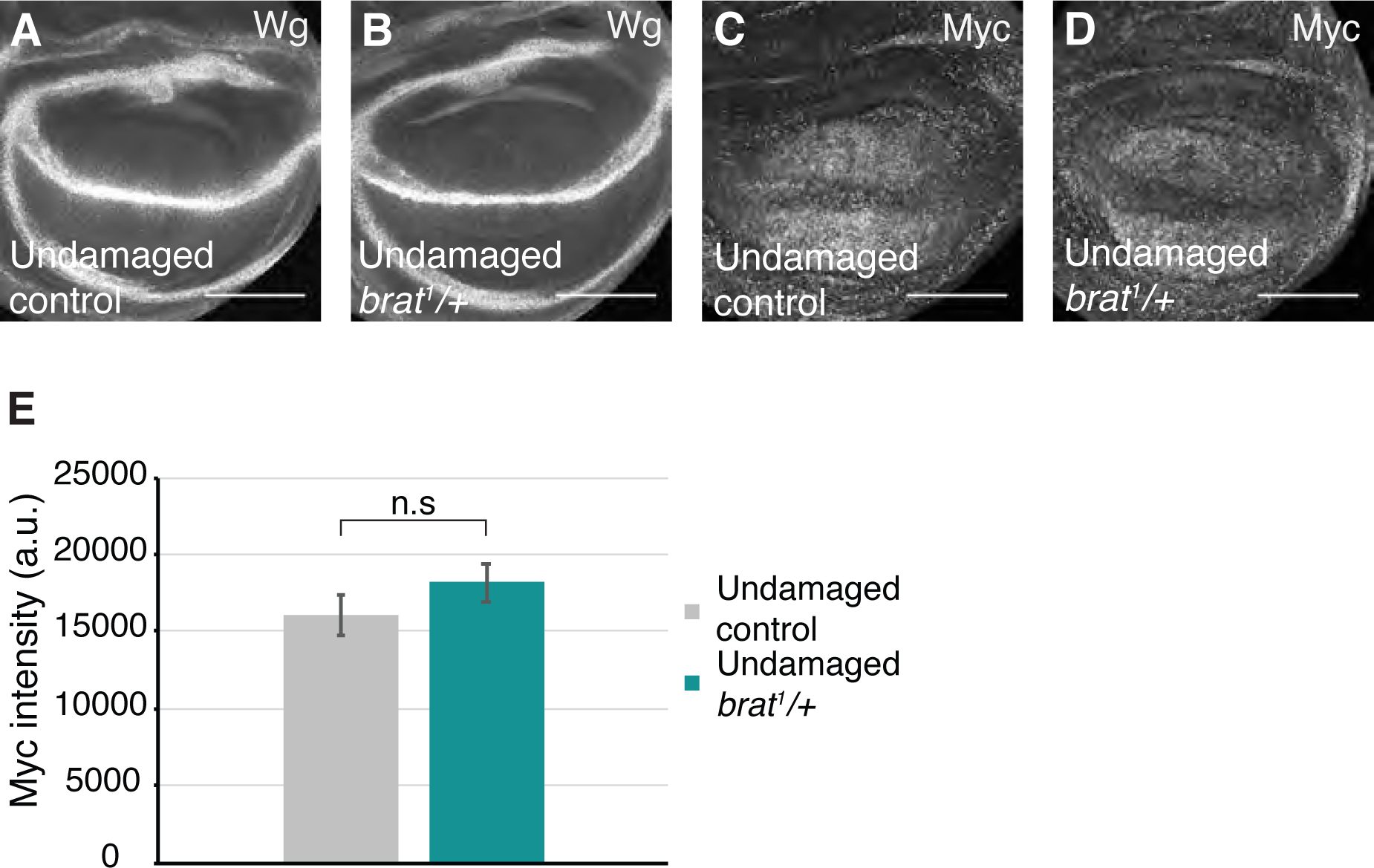
Effects of reduced Brat on Wg and Myc expression are regeneration specific (Related to. **Figure 3**). (A-B) Anti-Wg immunostaining in an undamaged control (*w^1118^*) disc (A) and an undamaged *brat^1^/+* disc (B). (C-D) Anti-Myc immunostaining in an undamaged control (*w^1118^*) disc (C) and an undamaged *brat^1^/+* disc (D). (E) Quantification of Myc fluorescence intensity in undamaged control (*w^1118^*) (n = 10) and *brat^1^/+* (n = 10) discs. Area for fluorescence intensity measurement was defined by wing pouch morphology and the elevated Myc expression domain in the wing pouch. Error bars represent SEM. Student’s T-test used for statistical analyses. Scale bars are 100 μm.

**Figure S4.**
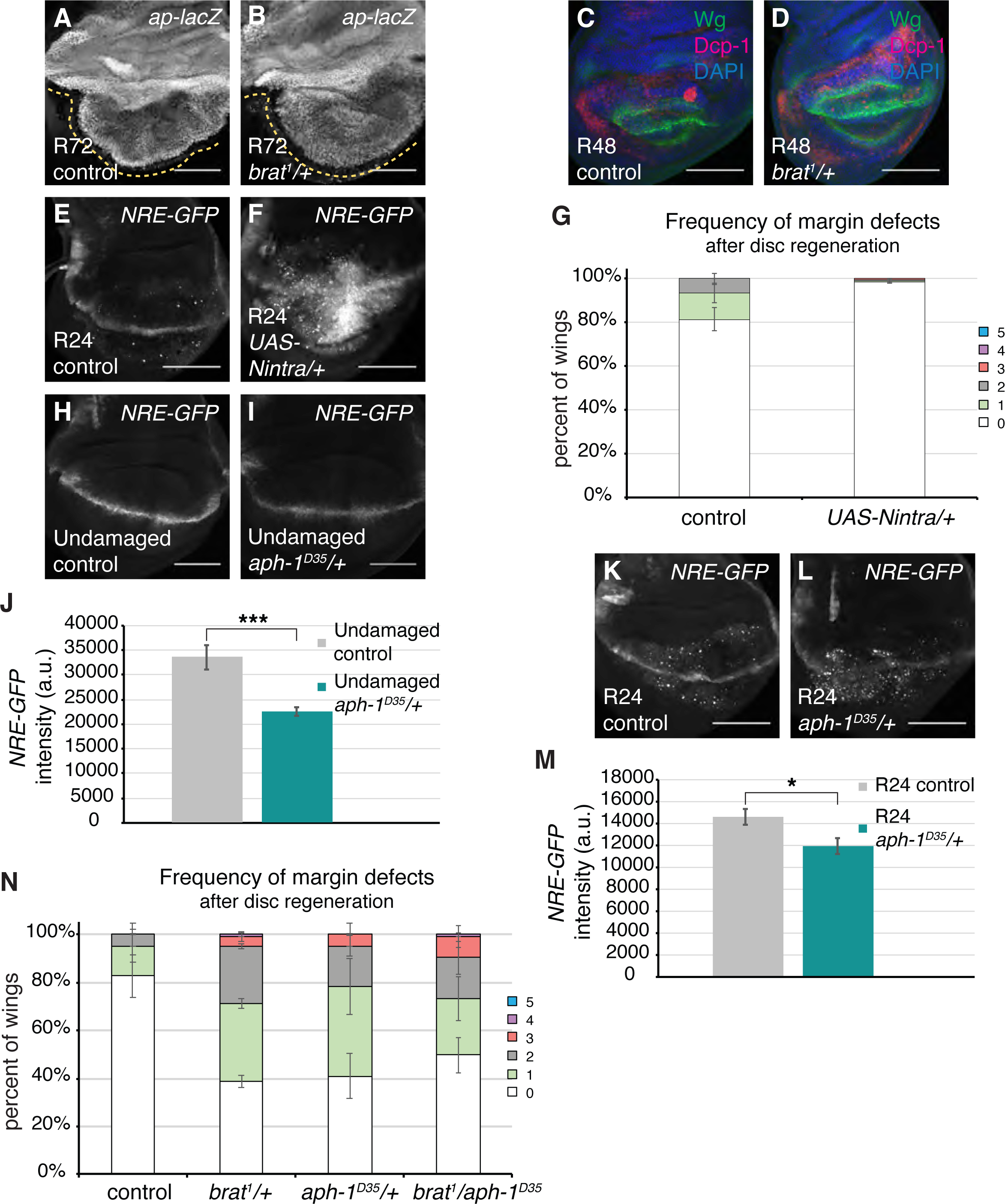
Elevated Notch signaling does not cause margin defects (Related to. **Figure 4**). (A-B) *ap-lacZ* expression in an R72 control (*w^1118^*) disc (A) and an R72 *brat^1^/+* disc (B). Dashed yellow lines are drawn next to the DV boundary to highlight it. (C-D) Anti-Wg (green) and anti-cleaved Dcp1 (red) immunostaining in an R48 control (*w^1118^*) disc (C) and an R48 *brat^1^/+* disc (D). (E-F) *NRE-GFP* expression in an R24 control (*w^1118^*) disc (E) and an R24 *UAS-Nintra/+* disc (F). (G) Frequency of margin defects seen in adult wings after disc regeneration for control (*w^1118^*) (n = 84) and *UAS-Nintra/+* (n = 357) wings, from five independent experiments. (H-I) *NRE-GFP* expression in an undamaged control (*w^1118^*) disc (H) and an undamaged *aph-1^D35^/+* disc (I). *NRE-GFP*/+ and *NRE- GFP*/*aph-1^D35^* animals were raised at room temperature and dissected during third instar. (J) Quantification of GFP intensity in undamaged control (*w^1118^*) (n = 15) and *aph- 1^D35^/+* (n = 15) discs. *** p < 0.0006. (K-L) *NRE-GFP* expression in an R24 control (*w^1118^*) disc (K) and an R24 *aph-1^D35^/+* disc (L). (M) Quantification of GFP intensity in R24 control (*w^1118^*) (n = 13) and R24 *aph-1^D35^/+* (n = 11) discs. * p < 0.02. (N) Frequency of margin defects in adult wings after disc regeneration for control (*w^1118^*) (n = 21), *brat^1^/+* (n = 137), *aph-1^D35^/+* (n = 38) and *brat^1^/aph-1^D35^*(n = 80) wings. Error bars represent SEM. Student’s T-test used for statistical analyses. Scale bars are 100 μm.

**Figure S5.**
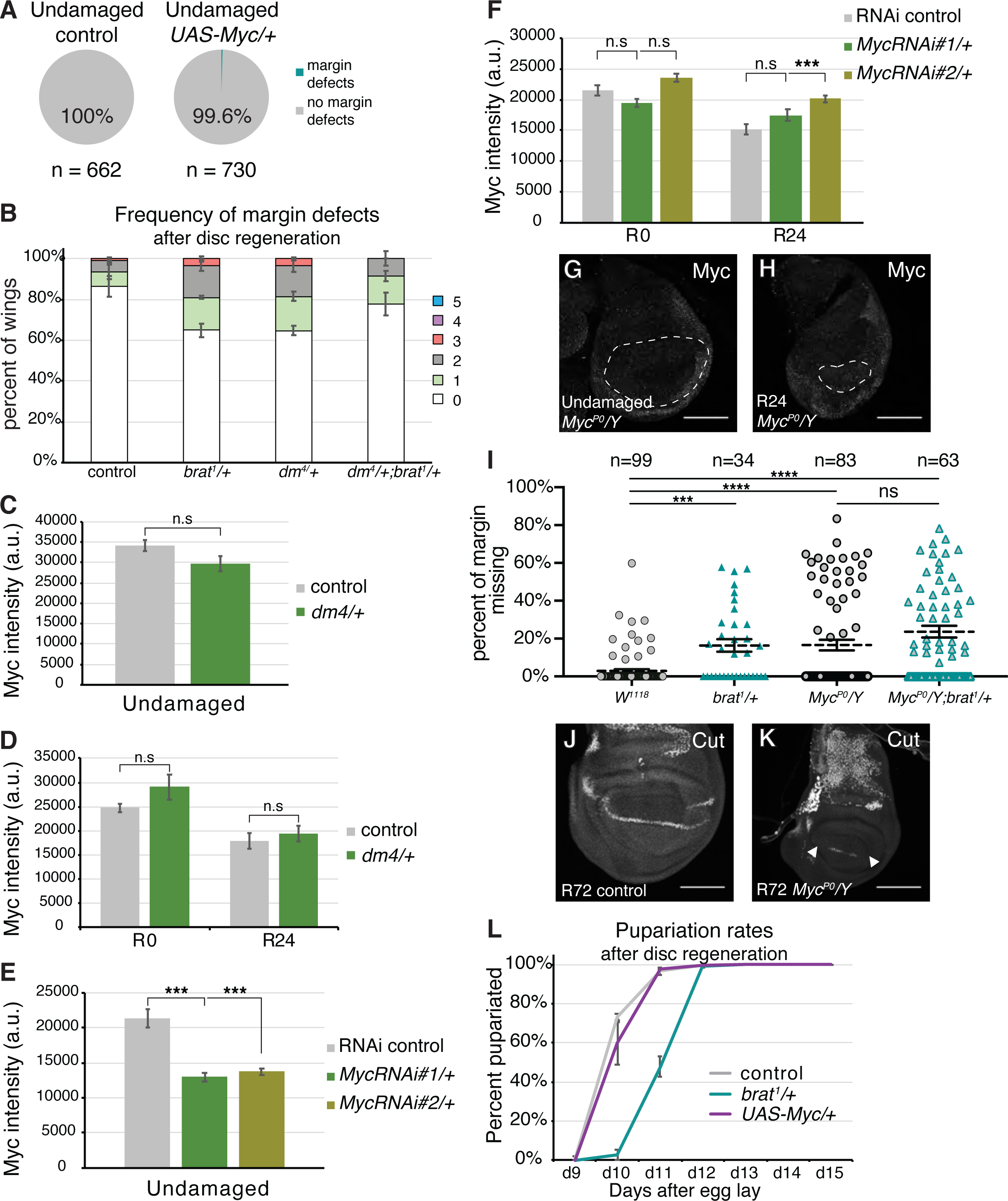
Reduction of Myc expression during regeneration also disrupts margin fate specification (Related to. **Figure 5**). (A) Margin defects detected in adult wings from undamaged control (*w^1118^*) and *UAS- Myc/+* discs. *rnGAL4, GAL80^ts^/TM6B* females were crossed to *w^1118^* or *UAS-Myc* males and taken through the protocol shown in Figure 1A. (B) Frequency of margin defects in adult wings after disc regeneration for control (*w^1118^*) (n = 103), *brat^1^/+* (n = 203), *dm^4^/+* (n = 94) and *dm^4^/+; brat^1^/+* (n = 94) wings, from three independent experiments. (C) Quantification of Myc fluorescence intensity in undamaged control (*w^1118^*) (n = 12) and *dm^4^/+* (n = 11) discs. *w^1118^* males were crossed to *w^1118^*or *dm^4^/FM7i, ActGFP* females and dissected when the animals were third instar. Area for fluorescence intensity measurement was defined by wing pouch morphology and the elevated Myc expression domain in the wing pouch. (D) Quantification of Myc fluorescence intensity in R0 control (*w^1118^*) (n = 13), R0 *dm^4^/+* (n = 10), R24 control (*w^1118^*) (n = 13), and R24 *dm^4^/+* (n = 10) discs. Area for fluorescence intensity measurement was defined by the elevated Myc expression domain in the wing pouch. (E) Quantification of Myc fluorescence intensity in undamaged control (VDRC genetic background line, called control) (n = 14), *MycRNAi#1/+* (n = 12), and *MycRNAi#2/+* (n = 13) discs. *rnGAL4, GAL80^ts^/TM6B* females were crossed to the control, *MycRNAi#1*, or *MycRNAi#2* males. The animals were shifted to 30°C during early third instar and kept there for 28 hours then dissected. *MycRNAi#1/+* *** p < 0.000007, *MycRNAi#2/+* *** p < 0.00002. Area for fluorescence intensity measurement was defined by wing pouch morphology. (F) Quantification of Myc fluorescence intensity in R0 control (n = 13), R0 *MycRNAi#1/+* (n = 15), R0 *MycRNAi#2/+* (n = 13), R24 control (n = 13), R24 *MycRNAi#1/+* (n = 13), and R24 *MycRNAi#2/+* (n = 13) discs. Fluorescence intensity was measured in the area marked by Anti-Nubbin immunostaining. *** p < 0.00007. (G,H) Anti-Myc immunostaining in undamaged (G) and regenerating R24 (H) *Myc^P0^/Y* imaginal wing discs. Dashed line outlines the wing pouch defined by anti-Nubbin immunostaining. (I) Margin tissue lost as a percentage of total wing perimeter for adult wings after disc damage and regeneration in the noted genotypes. (J, K) Immunostaining for Cut in control (J) and *Myc^P0^/Y* (K) regenerated (R72) discs showing margin disruption in the mutants.(L) Pupariation rates after disc regeneration for control (*w^1118^*) (n = 216), *brat^1^/+* (n = 114) and *UAS-Myc/+* (n = 209) animals, from three independent experiments. Error bars represent SEM. Student’s T-test used for statistical analyses.

**Figure S6.**
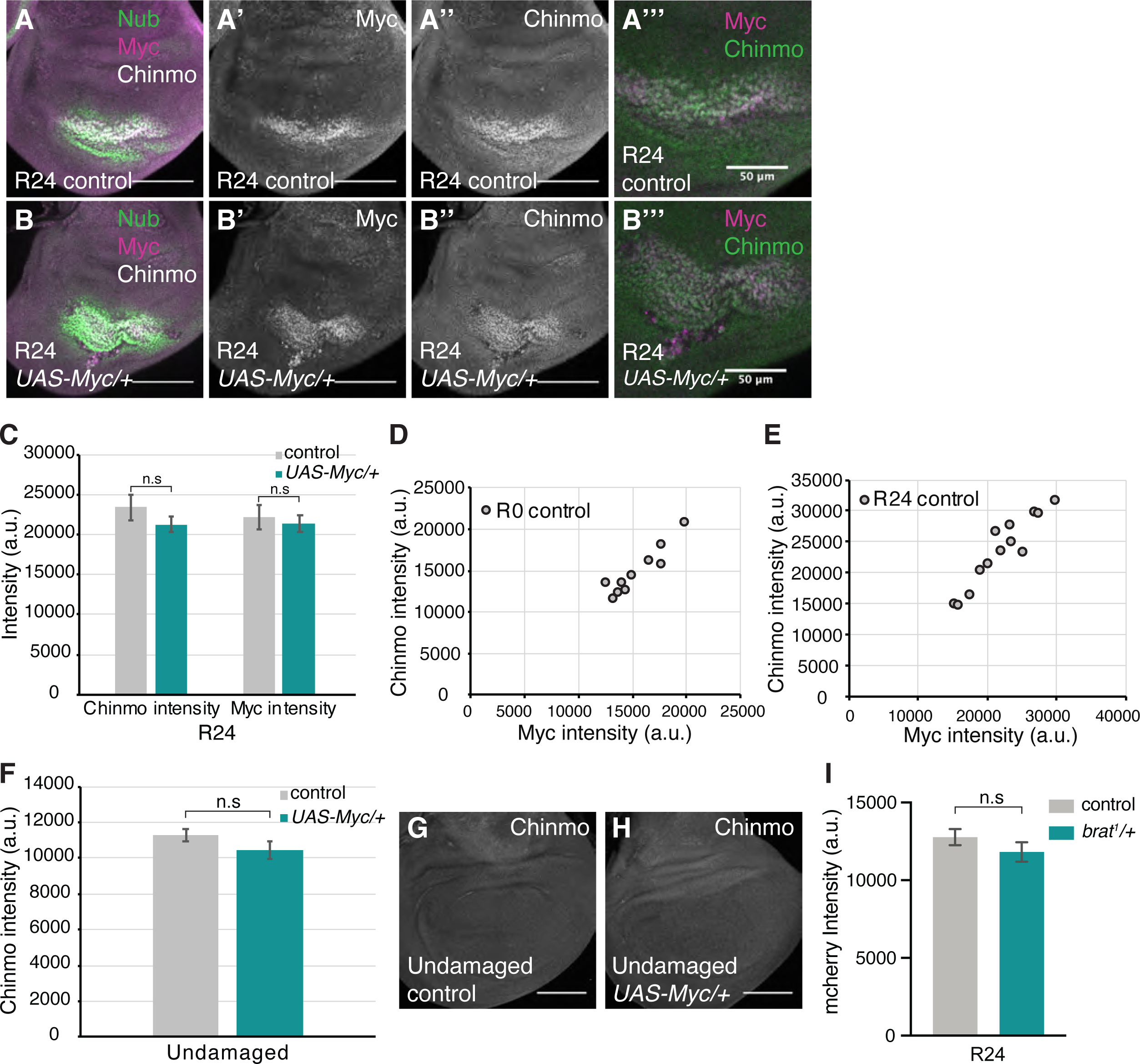
Myc regulates Chinmo expression (Related to. **Figure 7**). (A) Merge of anti-Nubbin, anti-Myc and anti-Chinmo immunostaining in an R24 control (*w^1118^*) disc. (A’-A’’) Same disc as (A) showing anti-Myc and anti-Chinmo immunostaining, respectively. (A’’’) Same disc as (A) showing an enlarged merge of anti-Myc and anti-Chinmo immunostaining. (B) Merge of anti-Nubbin, anti-Myc and anti- Chinmo immunostaining in an R24 *UAS-Myc/+* disc. (B’-B’’) Same disc as (B) showing anti-Myc and anti-Chinmo immunostaining, respectively. (B’’’) Same disc as (B) showing an enlarged merge of anti-Myc and anti-Chinmo immunostaining. (C) Quantification of Chinmo and Myc fluorescence intensity in R24 control (*w^1118^*) (n = 13) and R24 *UAS- Myc/+* (n = 14) discs. Area for fluorescence intensity measurement was defined by the elevated Myc expression domain in the wing pouch. Note that Myc and Chinmo expression co-localize. (D-E) Scatter plot showing correlation between Myc and Chinmo expression levels at R0 (D) and R24 (E). Pearson correlation coefficient for R0 = 0.93 and R24 = 0.94. (F) Quantification of Chinmo fluorescence intensity in undamaged discs dissected 31 hours after animals were shifted to 30°C on day 7 AEL. Control (*+ ; rnGAL4,GAL80ts/+*) (n = 14) and *UAS-Myc (UAS-Myc/+ ; rnGAL4, GAL80ts/+)* (n = 14). Area for fluorescence intensity measurement was defined by the Myc expression domain in the wing pouch. (G-H) Anti-Chinmo immunostaining in an undamaged control disc (G) and an undamaged *UAS-Myc/+* disc (H). (I) Quantification of fluorescence intensity from the *UAS-mCherry-chinmoUTR* transgene in control (n=16) and brat/+ (n=15) R24 regenerating discs. Area for fluorescence intensity measurement was defined by Nubbin expression. Error bars represent SEM. Student’s T-test used for statistical analyses. Scale bars are 100 μm, unless stated otherwise.

## References

1. Bosch M, Serras F, Martín-Blanco E, Baguñà J. JNK signaling pathway required for wound healing in regenerating Drosophila wing imaginal discs. Dev Biol. 2005;280(1):73–86.

2. Bergantiños C, Corominas M, Serras F. Cell death-induced regeneration in wing imaginal discs requires JNK signalling. Development. 2010;137(7):1169–79.

3. Bosch M, Bishop SA, Baguña J, Couso JP. Leg regeneration in Drosophila abridges the normal developmental program. Int J Dev Biol. 2010;54(8–9):1241–50.

4. Tasaki J, Shibata N, Sakurai T, Agata K, Umesono Y. Role of c-Jun N-terminal kinase activation in blastema formation during planarian regeneration. Dev Growth Differ. 2011;53(3):389–400.

5. Martín R, Pinal N, Morata G. Distinct regenerative potential of trunk and appendages of Drosophila mediated by JNK signalling. Development. 2017;144(21):dev.155507.

6. Bando T, Ishimaru Y, Kida T, Hamada Y, Matsuoka Y, Nakamura T, et al. Analysis of RNA-Seq data reveals involvement of JAK/STAT signalling during leg regeneration in the cricket Gryllus bimaculatus. Development. 2013;140(5):959–64.

7. Katsuyama T, Comoglio F, Seimiya M, Cabuy E, Paro R. During Drosophila disc regeneration, JAK/STAT coordinates cell proliferation with Dilp8-mediated developmental delay. Proc National Acad Sci. 2015;112(18):E2327–36.

8. Verghese S, Su TT. STAT, Wingless, and Nurf-38 determine the accuracy of regeneration after radiation damage in Drosophila. Plos Genet. 2017;13(10):e1007055.

9. Nakamura T, Mito T, Bando T, Ohuchi H, Noji S. Molecular and Cellular Basis of Regeneration and Tissue Repair. Cell Mol Life Sci. 2007;65(1):64.

10. Jiang H, Grenley MO, Bravo MJ, Blumhagen RZ, Edgar BA. EGFR/Ras/MAPK Signaling Mediates Adult Midgut Epithelial Homeostasis and Regeneration in Drosophila. Cell Stem Cell. 2011;8(1):84–95.

11. Fan Y, Wang S, Hernandez J, Yenigun VB, Hertlein G, Fogarty CE, et al. Genetic Models of Apoptosis-Induced Proliferation Decipher Activation of JNK and Identify a Requirement of EGFR Signaling for Tissue Regenerative Responses in Drosophila. Plos Genet. 2014;10(1):e1004131.

12. Jin Y, Ha N, Forés M, Xiang J, Gläßer C, Maldera J, et al. EGFR/Ras Signaling Controls Drosophila Intestinal Stem Cell Proliferation via Capicua-Regulated Genes. Plos Genet. 2015;11(12):e1005634.

13. Bando T, Mito T, Maeda Y, Nakamura T, Ito F, Watanabe T, et al. Regulation of leg size and shape by the Dachsous/Fat signalling pathway during regeneration. Development. 2009;136(13):2235–45.

14. Sun G, Irvine KD. Regulation of Hippo signaling by Jun kinase signaling during compensatory cell proliferation and regeneration, and in neoplastic tumors. Dev Biol. 2011;350(1):139–51.

15. Grusche FA, Degoutin JL, Richardson HE, Harvey KF. The Salvador/Warts/Hippo pathway controls regenerative tissue growth in Drosophila melanogaster. Dev Biol. 2011;350(2):255–66.

16. Hayashi S, Ochi H, Ogino H, Kawasumi A, Kamei Y, Tamura K, et al. Transcriptional regulators in the Hippo signaling pathway control organ growth in Xenopus tadpole tail regeneration. Dev Biol. 2014;396(1):31–41.

17. Grijalva JL, Huizenga M, Mueller K, Rodriguez S, Brazzo J, Camargo F, et al. Dynamic alterations in Hippo signaling pathway and YAP activation during liver regeneration. Am J Physiol-gastr L. 2014;307(2):G196–204.

18. Hobmayer B, Rentzsch F, Kuhn K, Happel CM, Laue CC von, Snyder P, et al. WNT signalling molecules act in axis formation in the diploblastic metazoan Hydra. Nature. 2000;407(6801):186.

19. Kawakami Y, Esteban CR, Raya M, Kawakami H, Martí M, Dubova I, et al. Wnt/β- catenin signaling regulates vertebrate limb regeneration. Gene Dev. 2006;20(23):3232– 7.

20. McClure KD, Schubiger G. A screen for genes that function in leg disc regeneration in Drosophila melanogaster. Mech Develop. 2008;125(1–2):67–80.

21. Schubiger M, Sustar A, Schubiger G. Regeneration and transdetermination: The role of wingless and its regulation. Dev Biol. 2010;347(2):315–24.

22. Wehner D, Cizelsky W, Vasudevaro MD, Özhan G, Haase C, Kagermeier-Schenk B, et al. Wnt/β-Catenin Signaling Defines Organizing Centers that Orchestrate Growth and Differentiation of the Regenerating Zebrafish Caudal Fin. Cell Reports. 2014;6(3):467–81.

23. Smith-Bolton RK, Worley MI, Kanda H, Hariharan IK. Regenerative Growth in Drosophila Imaginal Discs Is Regulated by Wingless and Myc. Dev Cell. 2009;16(6):797–809.

24. Hanovice NJ, Leach LL, Slater K, Gabriel AE, Romanovicz D, Shao E, et al. Regeneration of the zebrafish retinal pigment epithelium after widespread genetic ablation. Plos Genet. 2019;15(1):e1007939.

25. Harris RE, Setiawan L, Saul J, Hariharan IK. Localized epigenetic silencing of a damage-activated WNT enhancer limits regeneration in mature Drosophila imaginal discs. Elife. 2016;5:e11588.

26. Muneoka K, Bryant SV. Evidence that patterning mechanisms in developing and regenerating limbs are the same. Nature. 1982;298(5872):298369a0.

27. Gupta V, Gemberling M, Karra R, Rosenfeld GE, Evans T, Poss KD. An Injury- Responsive Gata4 Program Shapes the Zebrafish Cardiac Ventricle. Curr Biol. 2013;23(13):1221–7.

28. Roensch K, Tazaki A, Chara O, Tanaka EM. Progressive Specification Rather than Intercalation of Segments During Limb Regeneration. Science. 2013;342(6164):1375–9.

29. Mader MM, Cameron DA. Photoreceptor Differentiation during Retinal Development, Growth, and Regeneration in a Metamorphic Vertebrate. J Neurosci. 2004;24(50):11463–72.

30. McCusker CD, Gardiner DM. Positional Information Is Reprogrammed in Blastema Cells of the Regenerating Limb of the Axolotl (Ambystoma mexicanum). Plos One. 2013;8(9):e77064.

31. Myohara M. Differential tissue development during embryogenesis and regeneration in an annelid. Dev Dynam. 2004;231(2):349–58.

32. Schuster KJ, Smith-Bolton RK. Taranis Protects Regenerating Tissue from Fate Changes Induced by the Wound Response in Drosophila. Dev Cell. 2015;34(1):119–28.

33. Luttrell SM, Gotting K, Ross E, Alvarado AS, Swalla BJ. Head regeneration in hemichordates is not a strict recapitulation of development. Dev Dynam. 2016;245(12):1159–75.

34. Vizcaya-Molina E, Klein CC, Serras F, Mishra RK, Guigó R, Corominas M. Damage- responsive elements in Drosophila regeneration. Genome Res. 2018;28(12):1852–66.

35. Sun G, Irvine KD. Chapter Four Control of Growth During Regeneration. Curr Top Dev Biol. 2014;108:95–120.

36. Hariharan IK, Serras F. Imaginal disc regeneration takes flight. Curr Opin Cell Biol. 2017;48:10–6.

37. Sonoda J, Wharton RP. Drosophila Brain Tumor is a translational repressor. Gene Dev. 2001;15(6):762–73.

38. Loedige I, Stotz M, Qamar S, Kramer K, Hennig J, Schubert T, et al. The NHL domain of BRAT is an RNA-binding domain that directly contacts the hunchback mRNA for regulation. Gene Dev. 2014;28(7):749–64.

39. Loedige I, Jakob L, Treiber T, Ray D, Stotz M, Treiber N, et al. The Crystal Structure of the NHL Domain in Complex with RNA Reveals the Molecular Basis of Drosophila Brain-Tumor-Mediated Gene Regulation. Cell Reports. 2015;13(6):1206–20.

40. Arama E, Dickman D, Kimchie Z, Shearn A, Lev Z. Mutations in the β-propeller domain of the Drosophila brain tumor (brat) protein induce neoplasm in the larval brain. Oncogene. 2000;19(33):1203706.

41. Harris RE, Pargett M, Sutcliffe C, Umulis D, Ashe HL. Brat Promotes Stem Cell Differentiation via Control of a Bistable Switch that Restricts BMP Signaling. Dev Cell. 2011;20(1):72–83.

42. Betschinger J, Mechtler K, Knoblich JA. Asymmetric Segregation of the Tumor Suppressor Brat Regulates Self-Renewal in Drosophila Neural Stem Cells. Cell. 2006;124(6):1241–53.

43. Lee CY, Wilkinson BD, Siegrist SE, Wharton RP, Doe CQ. Brat Is a Miranda Cargo Protein that Promotes Neuronal Differentiation and Inhibits Neuroblast Self-Renewal. Dev Cell. 2006;10(4):441–9.

44. Chen G, Kong J, Tucker-Burden C, Anand M, Rong Y, Rahman F, et al. Human Brat Ortholog TRIM3 Is a Tumor Suppressor That Regulates Asymmetric Cell Division in Glioblastoma. Cancer Res. 2014;74(16):4536–48.

45. Schwamborn JC, Berezikov E, Knoblich JA. The TRIM-NHL Protein TRIM32 Activates MicroRNAs and Prevents Self-Renewal in Mouse Neural Progenitors. Cell. 2009;136(5):913–25.

46. Kudryashova E, Kramerova I, Spencer MJ. Satellite cell senescence underlies myopathy in a mouse model of limb-girdle muscular dystrophy 2H. J Clin Invest. 2012;122(5):1764–76.

47. Nicklas S, Otto A, Wu X, Miller P, Stelzer S, Wen Y, et al. TRIM32 Regulates Skeletal Muscle Stem Cell Differentiation and Is Necessary for Normal Adult Muscle Regeneration. Plos One. 2012;7(1):e30445.

48. Vonesch SC, Lamparter D, Mackay TFC, Bergmann S, Hafen E. Genome-Wide Analysis Reveals Novel Regulators of Growth in Drosophila melanogaster. Plos Genet. 2016;12(1):e1005616.

49. Skinner A, Khan SJ, Smith-Bolton RK. Trithorax regulates systemic signaling during Drosophila imaginal disc regeneration. Development. 2015;142(20):3500–11.

50. Khan SJ, Abidi SNF, Skinner A, Tian Y, Smith-Bolton RK. The Drosophila Duox maturation factor is a key component of a positive feedback loop that sustains regeneration signaling. Plos Genet. 2017;13(7):e1006937.

51. Brock AR, Seto M, Smith-Bolton RK. Cap-n-Collar Promotes Tissue Regeneration by Regulating ROS and JNK Signaling in the Drosophila melanogaster Wing Imaginal Disc. Genetics. 2017;206(3):1505–20.

52. Luschnig S, Moussian B, Krauss J, Desjeux I, Perkovic J, Nüsslein-Volhard C. An F1 Genetic Screen for Maternal-Effect Mutations Affecting Embryonic Pattern Formation in Drosophila melanogaster. Genetics. 2004;167(1):325–42.

53. Wright TRF, Beermann W, Marsh JL, Bishop CP, Steward R, Black BC, et al. The genetics of dopa decarboxylase in Drosophila melanogaster. Chromosoma. 1981;83(1):45–58.

54. Ferreira A, Boulan L, Perez L, Milán M. Mei-P26 Mediates Tissue-Specific Responses to the Brat Tumor Suppressor and the dMyc Proto-Oncogene in Drosophila. Genetics. 2014;198(1):249–58.

55. Halme A, Cheng M, Hariharan IK. Retinoids Regulate a Developmental Checkpoint for Tissue Regeneration in Drosophila. Curr Biol. 2010;20(5):458–63.

56. Colombani J, Andersen DS, Léopold P. Secreted Peptide Dilp8 Coordinates *Drosophila* Tissue Growth with Developmental Timing. Science. 2012;336(6081):582–5.

57. Garelli A, Gontijo AM, Miguela V, Caparros E, Dominguez M. Imaginal Discs Secrete Insulin-Like Peptide 8 to Mediate Plasticity of Growth and Maturation. Science. 2012;336(6081):579–82.

58. Abbott LC, Karpen GH, Schubiger G. Compartmental restrictions and blastema formation during pattern regulation in Drosophila imaginal leg discs. Dev Biol. 1981;87(1):64–75.

59. Ng M, Diaz-Benjumea FJ, Cohen SM. Nubbin encodes a POU-domain protein required for proximal-distal patterning in the Drosophila wing. Dev Camb Engl. 1995;121(2):589–99.

60. Komori H, Xiao Q, McCartney BM, Lee CY. Brain tumor specifies intermediate progenitor cell identity by attenuating β-catenin/Armadillo activity. Development. 2014;141(1):51–62.

61. Couso J, Bate M, Martinez-Arias A. A wingless-dependent polar coordinate system in Drosophila imaginal discs. Science. 1993;259(5094):484–9.

62. Wu DC, Johnston LA. Control of Wing Size and Proportions by Drosophila Myc. Genetics. 2010;184(1):199–211.

63. Diaz-Benjumea FJ, Cohen SM. Interaction between dorsal and ventral cells in the imaginal disc directs wing development in Drosophila. Cell. 1993;75(4):741–52.

64. Micchelli CA, Rulifson EJ, Blair SS. The function and regulation of cut expression on the wing margin of Drosophila: Notch, Wingless and a dominant negative role for Delta and Serrate. Dev Camb Engl. 1997;124(8):1485–95.

65. Becam I, Milán M. A permissive role of Notch in maintaining the DV affinity boundary of the Drosophila wing. Dev Biol. 2008;322(1):190–8.

66. Saj A, Arziman Z, Stempfle D, Belle W van Sauder U, Horn T, et al. A Combined Ex Vivo and In Vivo RNAi Screen for Notch Regulators in Drosophila Reveals an Extensive Notch Interaction Network. Dev Cell. 2010;18(5):862–76.

67. Mukherjee S, Tucker-Burden C, Zhang C, Moberg K, Read R, Hadjipanayis C, et al. Drosophila Brat and Human Ortholog TRIM3 Maintain Stem Cell Equilibrium and Suppress Brain Tumorigenesis by Attenuating Notch Nuclear Transport. Cancer Res. 2016;76(8):2443–52.

68. Couso JP, Bishop SA, Arias AM. The wingless signalling pathway and the patterning of the wing margin in Drosophila. Dev Camb Engl. 1994;120(3):621–36.

69. Jack J, Dorsett D, Delotto Y, Liu S. Expression of the cut locus in the Drosophila wing margin is required for cell type specification and is regulated by a distant enhancer. Dev Camb Engl. 1991;113(3):735–47.

70. Pierce SB, Yost C, Britton JS, Loo LWM, Flynn EM, Edgar BA, et al. dMyc is required for larval growth and endoreplication in Drosophila. Development. 2004;131(10):2317–27.

71. Bosch JA, Sumabat TM, Hariharan IK. Persistence of RNAi-Mediated Knockdown in Drosophila Complicates Mosaic Analysis Yet Enables Highly Sensitive Lineage Tracing. Genetics. 2016;203(1):109–18.

72. Johnston LA, Prober DA, Edgar BA, Eisenman RN, Gallant P. Drosophila myc Regulates Cellular Growth during Development. Cell. 1999;98(6):779–90.

73. Dillard C, Narbonne-Reveau K, Foppolo S, Lanet E, Maurange C. Two distinct mechanisms silence chinmo in Drosophila neuroblasts and neuroepithelial cells to limit their self-renewal. Development. 2017;145(2):dev.154534.

74. Flaherty MS, Salis P, Evans CJ, Ekas LA, Marouf A, Zavadil J, et al. chinmo Is a Functional Effector of the JAK/STAT Pathway that Regulates Eye Development, Tumor Formation, and Stem Cell Self-Renewal in Drosophila. Dev Cell. 2010;18(4):556–68.

75. Narbonne-Reveau K, Maurange C. Developmental regulation of regenerative potential in Drosophila by ecdysone through a bistable loop of ZBTB transcription factors. Plos Biol. 2019;17(2):e3000149.

76. Kudron MM, Victorsen A, Gevirtzman L, Hillier LW, Fisher WW, Vafeados D, et al. The modERN Resource: Genome-Wide Binding Profiles for Hundreds of Drosophila and Caenorhabditis elegans Transcription Factors. Genetics. 2017;208(3):genetics.300657.2017.

77. Yang J, Sung E, Donlin-Asp PG, Corces VG. A subset of Drosophila Myc sites remain associated with mitotic chromosomes co-localized with insulator proteins. Nat Commun. 2013;4(1):1464–1464.

78. Sitaram P, Lu S, Harsh S, Herrera SC, Bach EA. Next-Generation Sequencing Reveals Increased Anti-oxidant Response and Ecdysone Signaling in STAT Supercompetitors in Drosophila. G3 Genes Genomes Genetics. 2019;9(8):2609–22.

79. Narbonne-Reveau K, Maurange C. Developmental regulation of regenerative potential in Drosophila by ecdysone through a bistable loop of ZBTB transcription factors. Plos Biol. 2019;17(2):e3000149.

80. Amati B, Land H. Myc—Max—Mad: a transcription factor network controlling cell cycle progression, differentiation and death. Curr Opin Genet Dev. 1994;4(1):102–8.

81. Takahashi K, Yamanaka S. Induction of Pluripotent Stem Cells from Mouse Embryonic and Adult Fibroblast Cultures by Defined Factors. Cell. 2006;126(4):663–76.

82. Marygold SJ, Coelho CMA, Leevers SJ. Genetic Analysis of RpL38 and RpL5, Two Minute Genes Located in the Centric Heterochromatin of Chromosome 2 of Drosophila melanogaster. Genetics. 2005;169(2):683–95.

83. Tian Y, Smith-Bolton RK. Regulation of growth and cell fate during tissue regeneration by the two SWI/SNF chromatin-remodeling complexes of Drosophila. Genetics. 2020;217(1):iyaa028.

84. Hayashi R, Goto Y, Ikeda R, Yokoyama KK, Yoshida K. CDCA4 Is an E2F Transcription Factor Family-induced Nuclear Factor That Regulates E2F-dependent Transcriptional Activation and Cell Proliferation. J Biol Chem. 2006;281(47):35633–48.

85. Hsu SI, Yang CM, Sim KG, Hentschel DM, O’Leary E, Bonventre JV. TRIP-Br: a novel family of PHD zinc finger- and bromodomain-interacting proteins that regulate the transcriptional activity of E2F-1/DP-1. Embo J. 2001;20(9):2273–85.

86. Watanabe-Fukunaga R, Iida S, Shimizu Y, Nagata S, Fukunaga R. SEI family of nuclear factors regulates p53-dependent transcriptional activation. Genes Cells. 2005;10(8):851–60.

87. Calgaro S, Boube M, Cribbs DL, Bourbon HM. The Drosophila gene taranis encodes a novel trithorax group member potentially linked to the cell cycle regulatory apparatus. Genetics. 2002;160(2):547–60.

88. Laver JD, Li X, Ray D, Cook KB, Hahn NA, Nabeel-Shah S, et al. Brain tumor is a sequence-specific RNA-binding protein that directs maternal mRNA clearance during the Drosophila maternal-to-zygotic transition. Genome Biol. 2015;16(1):94.

89. Komori H, Golden KL, Kobayashi T, Kageyama R, Lee CY. Multilayered gene control drives timely exit from the stem cell state in uncommitted progenitors during Drosophila asymmetric neural stem cell division. Gene Dev. 2018;32(23–24):1550–61.

90. Hazelrigg T, Levis R, Rubin GM. Transformation of white locus DNA in Drosophila: Dosage compensation, zeste interaction, and position effects. Cell. 1984;36(2):469–81.

91. Wright TR, Bewley GC, Sherald AF. The genetics of dopa decarboxylase in Drosophila melanogaster. II. Isolation and characterization of dopa-decarboxylase-deficient mutants and their relationship to the alpha-methyl-dopa-hypersensitive mutants. Genetics. 1976;84(2):287–310.

92. Parks AL, Cook KR, Belvin M, Dompe NA, Fawcett R, Huppert K, et al. Systematic generation of high-resolution deletion coverage of the Drosophila melanogaster genome. Nat Genet. 2004;36(3):ng1312.

93. Stathakis DG, Pentz ES, Freeman ME, Kullman J, Hankins GR, Pearlson NJ, et al. The genetic and molecular organization of the Dopa decarboxylase gene cluster of Drosophila melanogaster. Genetics. 1995;141(2):629–55.

94. Cohen B, McGuffin ME, Pfeifle C, Segal D, Cohen SM. apterous, a gene required for imaginal disc development in Drosophila encodes a member of the LIM family of developmental regulatory proteins. Gene Dev. 1992;6(5):715–29.

95. Littleton JT, Bellen HJ. Genetic and phenotypic analysis of thirteen essential genes in cytological interval 22F1-2; 23B1-2 reveals novel genes required for neural development in Drosophila. Genetics. 1994;138(1):111–23.

96. Averof M, Cohen SM. Evolutionary origin of insect wings from ancestral gills. Nature. 1997;385(6617):385627a0.

97. Classen AK, Bunker BD, Harvey KF, Vaccari T, Bilder D. A tumor suppressor activity of Drosophila Polycomb genes mediated by JAK-STAT signaling. Nat Genet. 2009;41(10):1150–5.

98. Mitchell NC, Johanson TM, Cranna NJ, Er ALJ, Richardson HE, Hannan RD, et al. Hfp inhibits Drosophila myc transcription and cell growth in a TFIIH/Hay-dependent manner. Development. 2010;137(17):2875–84.

